# Excess calorie intake early in life increases susceptibility to colitis in the adult

**DOI:** 10.1101/681759

**Authors:** Ziad Al Nabhani, Sophie Dulauroy, Emelyne Lécuyer, Bernadette Polomack, Marion Berard, Gérard Eberl

## Abstract

Epidemiological data report an association between obesity and inflammatory bowel disease (IBD) ^1–3^. Furthermore, animal models demonstrate that maternal high fat diet (HFD) and maternal obesity increase susceptibility to IBD in the offsprings ^4–8^. However, the mechanisms that translate maternal obesity and HFD into increased susceptibility to IBD later in life remain unknown. Here we report that excess calorie intake by neonatal mice, as a consequence of maternal HFD, forced feeding of neonates or low litter competition, lead to an increase, during weaning, in intestinal permeability, expression of pro-inflammatory cytokines and hydrogen sulfide production by the microbiota. In this context, intestinal permeability, cytokine expression and hydrogen sulfide engaged in a mutual positive feedback that imprinted increased susceptibility to colitis in the adult. This pathological imprinting was prevented by the neutralization of IFNγ and TNFα, of the production of hydrogen sulphide, or by normalization of intestinal permeability during weaning. Thus, excessive calorie intake by neonates leads to multiple causally-linked perturbations in the intestine that imprint the individual with long term susceptibility to IBD.

## Main Text

In order to assess the impact of excess calorie intake early in life on the susceptibility to colitis later in life, we used three distinct models of neonatal overfeeding that lead to overweight during weaning. Feeding mothers were given HFD (Fig. 1a), neonates were gavaged 2 times per day with a preparation of coconut oil (Fig. 1b), or litters were reduced to 3 pups while control litters included 7 pups (and thus benefited from less access to maternal milk) (Fig. 1c). All three conditions lead to overweight of the pups (Fig. 1d-f) characterized by increased fat depots (Supplementary Fig. 1a-c). The early overweight was transient and all groups of mice developed normal body weight at adult age (Supplementary Fig. 1d-f). However, mice that had experienced neonatal overweight showed increased susceptibility (or pathological imprinting) to dextran sodium sulfate (DSS)-induced colitis in adulthood, characterized by increased weight loss, higher disease activity index, increased intestinal permeability and inflammation, colon shortening and eventually, decreased survival (Fig. 1g-r and Supplementary Fig. 2-4).

**Figure 1.**
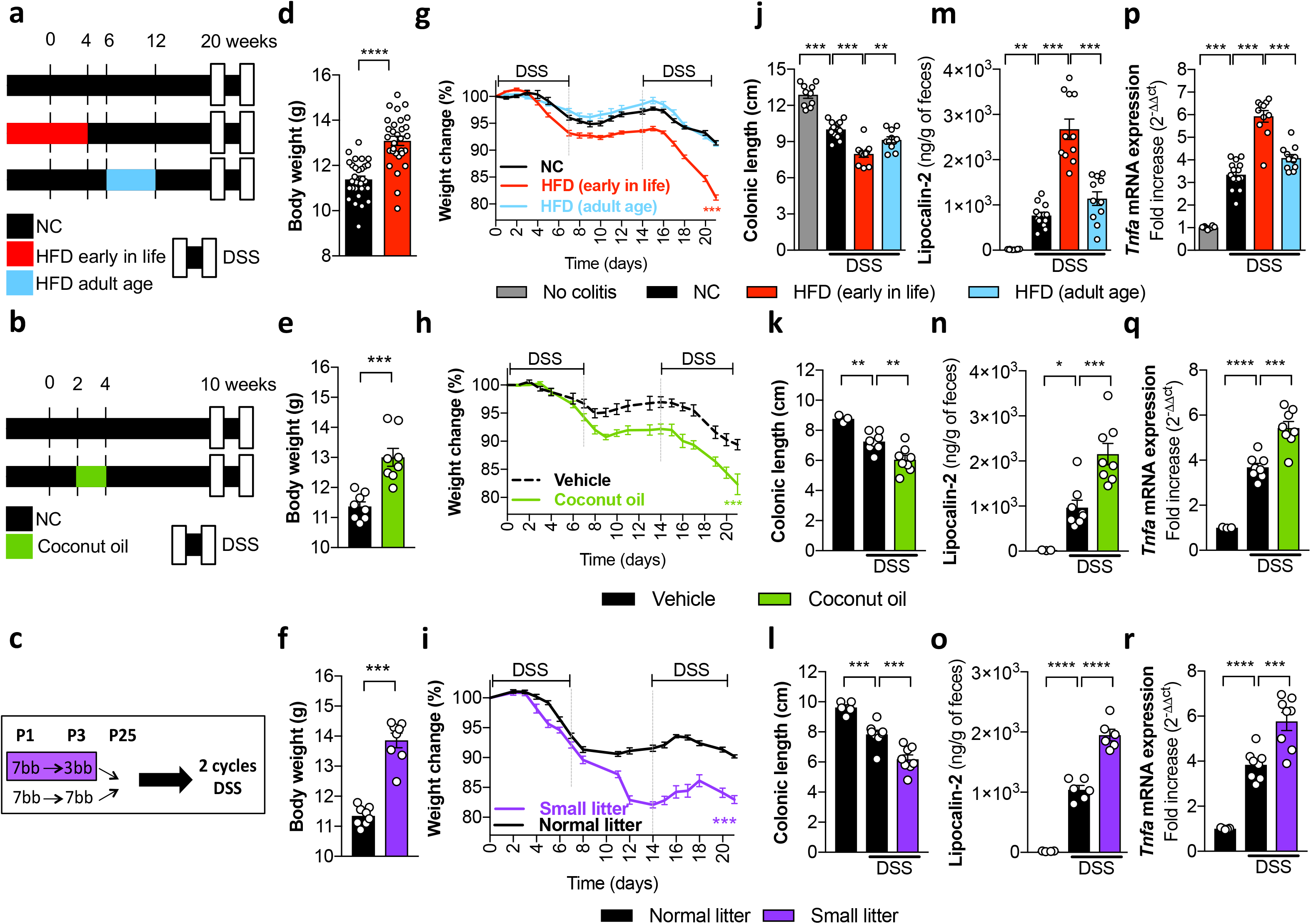
Excessive calorie intake early in life increases susceptibility to colitis in the adult. **(a)** Mice exposed to high fat diet (HFD) early in life were weaned to normal chow (NC) at 4 weeks of age, and mice fed HFD at adult age (between 6 and 12 weeks of age) were returned to NC. All group were challenged with 2 cycles of DSS at 20 weeks of age. **(b)** Mice fed NC were supplemented or not with coconut oil between 2 to 4 weeks after birth and then challenged with 2 cycles of DSS at 10 weeks of age. **(c)** The number of mice reared to NC was reduced at day 3 post-birth to 3 pups (SL; small litter) or maintained to 7 pups per litter as normal litter (NL) and then challenged with 2 cycles of DSS at 12 weeks of age. **(d-f)** Body weight measured at 4 weeks of age **(d)** in mice fed NC or HFD (n=30/group), **(e)** supplemented or not with coconut oil (n=8/group), and **(f)** grown in NL or SL (n=8/group). For **d-f** Mann Whitney test (\*\*\**P* < 0.001 and \*\*\*\**P* < 0.0001). **(g-i)** Percentage of body weight loss during 3 weeks after initiation of DSS **(g)** in mice fed NC (n=7), HFD early in life (n=5) or HFD at adult age (n=10), **(h)** supplemented or not with coconut oil (n = 8-9/group), or **(i)** grown in NL or SL (n=8/group). **(j-l)** Colonic length, **(m-o)** lipocalin-2 level, and **(p-r)** colonic expression of *Tnfa* mRNA measured at day 21 after initiation of DSS challenge **(j,m,p)** in mice fed NC, or HFD early in life or HFD at adult age (n=10-15/group), **(k,n,q)** supplemented or not with coconut oil (n=3-8/group), or **(l,o,r)** grown in NL or SL (n=5-8/group). One-way ANOVA with Sidak’s multiple comparisons test (\**P* < 0.05; \*\**P* < 0.01; \*\*\**P* < 0.001 and \*\*\*\**P* < 0.0001). ns = not significant. All data were pooled from at least two (d-r) independent experiments. All data are shown as mean ± s.e.m.

We next examined the effect of the three regimens on the inflammatory status and permeability of the intestine. All three regimens lead to an increased expression, at weaning but not at later time points (Supplementary Fig. 5), of transcripts coding for the pro-inflammatory cytokines TNFα, IFNγ, IL-1β, IL-12, IL-6 and IL-22, as well as a decrease in transcripts for the anti-inflammatory cytokine IL-10 (Fig. 2a-c and Supplementary Fig. 6). This was paralleled by an increase in intestinal permeability, as measured by serum levels of orally fed FITC-dextran (Fig. 2d-f), higher expression of *Mylk*, coding for the myosin light chain kinase MLCK that regulates the gut epithelial tight junctions, and lower expression of *Tjp1* and *Tjp2* coding for the tight junction proteins zona occludens 1 (ZO1) and ZO2 (Fig. 2a-c). The expression of other genes, involved in epithelial repair and mucus, such as *Tff3* and *Muc2*, were also affected (Supplementary Fig. 6). Abnormalities observed early in life were dependent on intestinal microbiota as most effects on the inflammatory reaction and increased permeability induced by neonatal overweight were prevented by concomitant treatment of the feeding mothers with a cocktail of antibiotics (Fig. 2 and Supplementary Fig. 6 and 7).

**Figure 2.**
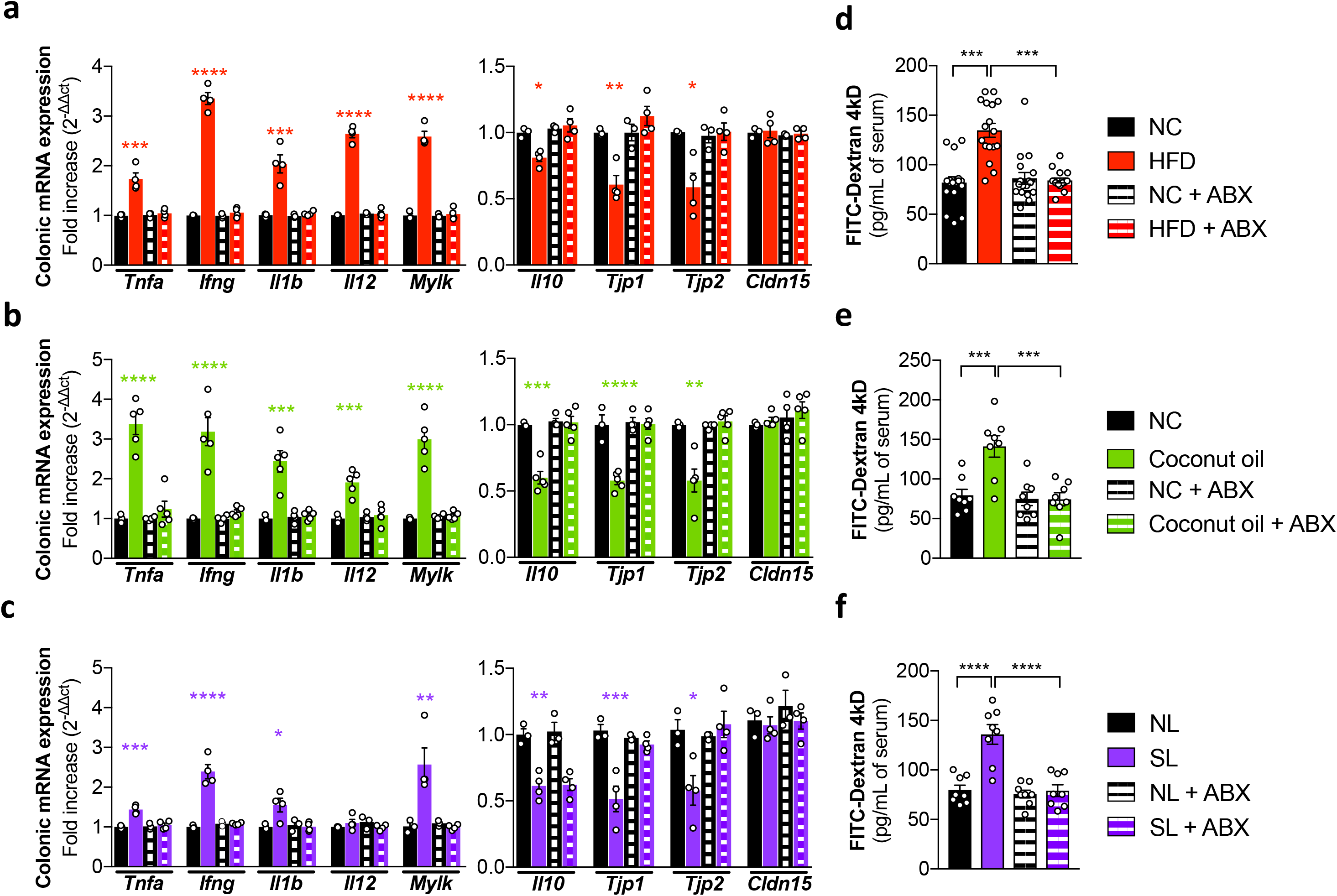
Excessive calorie intake early in life increases intestinal permeability and inflammation at weaning. **(a-c)** Colonic mRNA expression of the indicated genes, and **(d-f)** FITC-dextran 4kD level measured at 4 weeks of age **(a,d)** in mice fed NC or HFD, **(b,e)** supplemented or not with coconut oil, and **(c,f)** grown in NL or SL. One-way ANOVA with Sidak’s multiple comparisons test (\**P* < 0.05; \*\**P* < 0.01; \*\*\**P* < 0.001 and \*\*\*\**P* < 0.0001). For **a** (n=3-4/group)**, b** (n=4-6/group)**, c** (n=3-4/group), the data is representative of one among two independent experiments. For **d** (n=17/group)**, e** (n=6/group)**, f** (n=8/group), the data were pooled from at least two independent experiments. Data are shown as mean ± s.e.m.

We addressed whether the increased inflammation and intestinal permeability in overweight weaning mice induced increased susceptibility to colitis (or pathological imprinting) in adults. Offsprings of mothers fed HFD after birth were treated with neutralizing antibodies to IFNγ and TNFα from 2 to 4 weeks after birth. Inhibition of these two pro-inflammatory cytokines was sufficient to prevent overweight in weaning mice (Fig. 3a), to prevent the increase in expression of other pro-inflammatory cytokines (Fig. 3b), and normalized intestinal permeability (Fig. 3c and Supplementary Fig. 8). Most importantly, neutralization of IFNγ and TNFα during weaning also prevented increased susceptibility to DSS-induced colitis in adulthood (Fig. 3d-g and Supplementary Fig. 9). Similar results were obtained when MLCK was inhibited from 2 to 4 weeks after birth by the synthetic naphthalenesulphonyl derivative ML-7 (Fig. 3h-n and Supplementary Fig. 10,11). Blocking MLCK normalized intestinal permeability as expected, but also prevented the increase in the expression of pro-inflammatory cytokines, and, as a consequence, prevented pathological imprinting.

**Figure 3.**
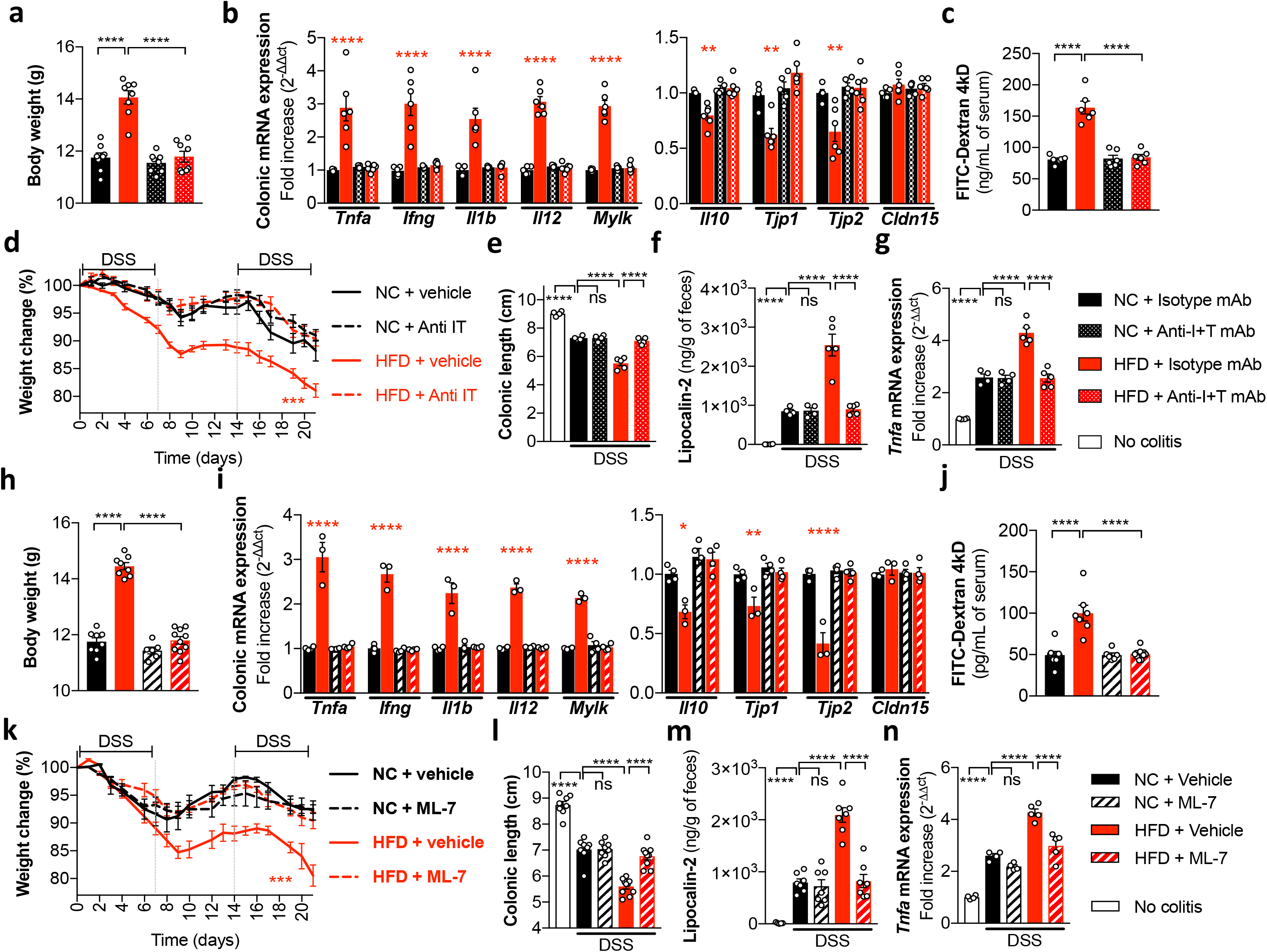
Prevention of increased adult susceptibility through gut normalization at weaning. **(a-g)** Mice fed NC or HFD early in life were treated with anti-TNFα and anti-IFNγ neutralizing antibodies during 2 to 4 weeks of age and then challenged with 2 cycles of DSS at 12 weeks of age. **(a)** Body weight (n=8/group), **(b)** transcript profile (n=5-6/group), and **(c)** intestinal permeability (n=6/group) were measured at 4 weeks of age. **(d)** Percentage of weight loss during 3 weeks after initiation of DSS challenge (n=6-7/group). **(e)** Colonic length **(f)** lipocalin-2 level, and **(g)** colonic expression of *Tnfa* mRNA measured at day 21 after initiation of DSS challenge (n=4-5/group). For **a, c, d, e, f, g**, the data were pooled from at least two independent experiments. For **b**, the data is representative of one among two independent experiments. **(h-n)** Mice fed NC or HFD early in life were treated with ML-7 (an inhibitor of MLCK) during 2 to 4 weeks of age and then challenged with 2 cycles of DSS at 12 weeks of age. **(h)** Body weight, **(i)** transcript profile, and **(j)** intestinal permeability were measured at 4 weeks of age. **(k)** Percentage of weight loss during 3 weeks after initiation of DSS challenge. **(l)** Colonic length **(m)** lipocalin-2 level, and **(n)** colonic expression of *Tnfa* mRNA measured at day 21 after initiation of DSS challenge. One-way ANOVA with Sidak’s multiple comparisons test (\**P* < 0.05; \*\**P* < 0.01; \*\*\**P* < 0.001 and \*\*\*\**P* < 0.0001). ns = not significant. For **h** (n=6-10/group)**, j** (n=6-10/group)**, k** (n=4-5/group)**, l** (n=8-10/group)**, m** (n=5-7/group)**, n** (n=4-5/group), the data were pooled from at least two independent experiments. For **i**(n=3-4/group), the data is representative of one among two independent experiments. All data are shown as mean ± s.e.m.

We next explored the mechanisms by which excessive calorie intake in neonates affected the expression of cytokines and gut permeability. Previous work shows that HFD favours the expansion of sulfide-producing bacteria ^9,10^. Hydrogen sulfide (H_2_S) leads to destabilization of the mucus layer by reducing S-S bonds in the protein network formed by essential mucus components such as Muc2 ^11,12^ and is toxic to colonic epithelial cells ^13^. In offsprings of mothers fed HFD, in neonates gavaged with coconut oil diet, and in mice raised in small litters, the expression of bacterial *dsrA* coding for the dissimilative sulfide reductase, a bacterial enzyme essential for the production of sulfide ^14^, was significantly increased during weaning, as was the concentration in fecal H_2_S (Fig. 4a-c and Supplementary Fig. 12a-f). This increase of in fecal H_2_S was prevented by the concomitant treatment of feeding mothers with a cocktail of antibiotics (Supplementary Fig. 12g,h). Blocking the production of H_2_S by bacteria with 5-aminosalicylic acid (5-ASA)^15^ (Supplementary Fig. 12i,j) prevented overweight in weaning mice, reduced the expression of several pro-inflammatory cytokines and normalized intestinal permeability (Fig. 4d-f and Supplementary Fig. 13). As a consequence, pathological imprinting was prevented, and the susceptibility of mice to DSS-induced colitis was reduced in adulthood (Fig. 4g and Supplementary Fig. 14). Furthermore, the gavage of GF mice from 2 to 4 weeks after birth with sodium hydrogen sulfide (NaHS) as rapid donor of H_2_S, in the presence of microbial immunogens (given as heat-killed microbiota), was sufficient to induce overweight when grown in small litters, to induce an increase in the expression of pro-inflammatory cytokines and an increase in intestinal permeability (Fig. 4i-k and Supplementary Fig. 15). Such mice developed pathological imprinting and increased susceptibility to DSS-induced colitis in adulthood (Fig. 4l,m and Supplementary Fig. 16). Similar results were obtained using the slow hydrogen sulfide-releasing compound GYY4137 ^16^ (Supplementary Fig. 17,18). Finally, pathological imprinting induced by NaHS gavage at weaning was prevented by the concomitant neutralization of TNFα and IFNγ, or by inhibition of MLCK (Supplementary Fig. 19).

**Figure 4.**
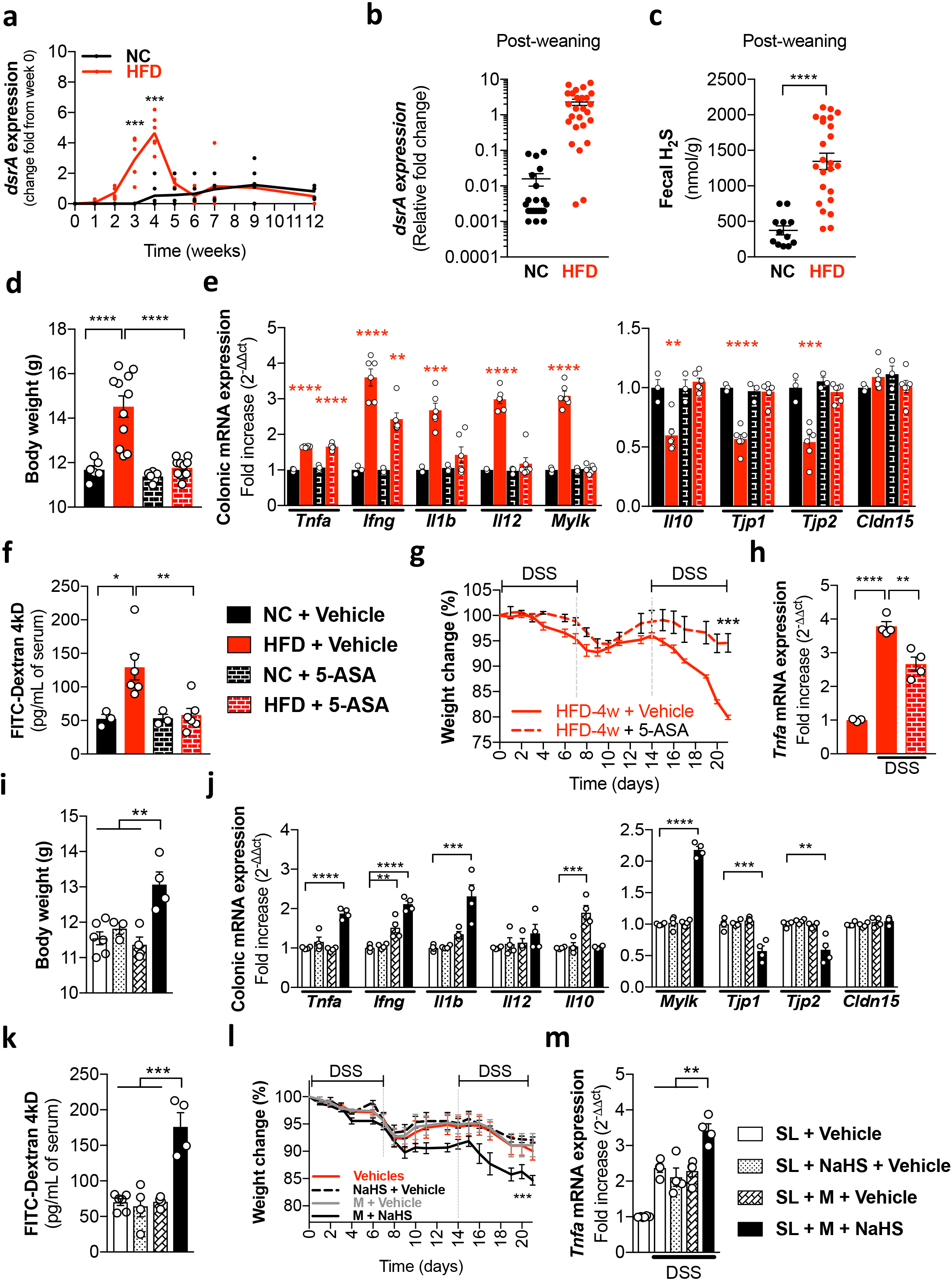
Involvement of H_2_S in pathological imprinting by excessive calorie intake early in life. **(a-b)** The expression of bacterial *dsrA* coding for the bacterial dissimilative sulfite reductase and **(b)** measured at 4 weeks after birth in mice fed NC or HFD. **(c)** Fecal H_2_S levels in mice fed HFD and control mice at 4 weeks of age. **(d-f)** Mice fed NC or HFD early in life are treated with 5-aminosalicylic acid (5-ASA) between 2 to 4 weeks after birth. **(d)** Body weight, **(e)** transcript profile and **(f)** intestinal permeability measured at 4 weeks after birth. **(g-h)** Mice fed HFD early in life were treated with 5-ASA between 2 to 4 weeks after birth and challenged with 2 cycles of DSS at 10 weeks of age. **(g)** Percentage of weight loss during 3 weeks after initiation of DSS challenge and **(h)** colonic expression of *Tnfa* mRNA measured at day 21 after initiation of DSS challenge. **(i-m)** Small litter of germ-free mice exposed to heat-killed microbiota (M) and/or to sodium hydrogen sulfide (NaHS) between 2 to 4 weeks after birth and then challenged with 2 cycles of DSS at 12 weeks of age. **(i)** Body weight, **(j)** transcript profile and **(k)** intestinal permeability measured at 4 weeks after birth. **(l)** Percentage of weight loss during 3 weeks after initiation of DSS challenge and **(m)** colonic expression of *Tnfa* mRNA measured at day 21 after initiation of DSS challenge. For **a, b, c** and **g** Mann Whitney test (\*\*\**P* < 0.001 and \*\*\*\**P* < 0.0001). For **d, e, f, h** and **i-m** one-way ANOVA with Sidak’s multiple comparisons test (\**P* < 0.05; \*\**P* < 0.01; \*\*\**P* < 0.001 and \*\*\*\**P* < 0.0001). ns = not significant. For **a** (n=6/group)**, b** (n=21-26/group)**, c** (n=12-24/group)**, d** (n=6-11/group), the data were pooled from at least two independent experiments. For **e** and **f** (n=3-6/group)**, g** and **h** (n=3-4/group), **i, j, l**, **m** (n=4-6/group)**, k** (n=4-6/group), the data is representative of one among two independent experiments. All data are shown as mean ± s.e.m.

Taken together, these results show that neonatal overweight induces pathological imprinting for increased susceptibility to colitis later in life. We report that overweight induced by HFD in feeding mothers, direct gavage of neonates with lipid-rich food or low littermate competition (and thus more access to maternal milk) lead to increased production of sulfide by the microbiota, increased intestinal permeability and increased expression of pro-inflammatory cytokines. These perturbations were causally-linked, and even if limited in time to the weaning period, lead to increased susceptibility to DSS-induced colitis later in life. Thus, the weaning period appears to be a critical time period for pathological imprinting, as exposure to HFD after weaning did not lead to increased susceptibility to colitis (Fig. 1g,j,m,p and Supplementary Fig. 2). Previous reports point to the notion that the time period between birth and weaning is a unique time window of opportunity for the normal development of the immune system, with pathological consequences later in life if perturbed ^17–22^. In the context of neonatal overweight, it remains to be assessed whether any type of excess calorie intake, through lipids, carbohydrates or proteins, induces pathological imprinting.

The intestinal and lung microbiota play an important role during this period, as mice not exposed to microbiota before weaning develop increased susceptibility to allergy and colitis in adulthood ^17–21^. A similar phenomenon is observed in human, as neonates treated with antibiotics develop increased susceptibility to allergy and pediatric colitis ^23,24^, a phenomenon framed by the hygiene hypothesis (stating that reduced exposure to microbes increases susceptibility to inflammatory pathologies) ^25^. In the context of neonatal overfeeding, we show that microbiota is a necessary element in the perturbation of intestinal permeability and inflammation, involving the production of hydrogen sulfide. Several bacterial taxa produce hydrogen sulfide in mice fed HFD ^9,10,15^. It remains to be understood how excessive calorie intake before and during weaning promotes the expansion of sulfide-producing bacteria, and, for preventive measures, how the expansion of such bacteria can be blocked.

The mechanisms by which neonatal perturbations translate into pathological imprinting remain poorly understood. In a model of intestinal allergic reaction induced by invariant NKT cells, microbiota induces modification, in dendritic cells, of the epigenetic code of *Cxcl16* involved in the recruitment of NKT cells ^17^. In a transgenic T cell receptor model of colitis, the induction, during weaning, of regulatory T cells carrying the transgenic TCR is required to prevent severe colitis in the adult ^22^. Finally, maternal microbiota shaped by antibiotic treatment can transfer increased susceptibility to colitis by vertical transmission to offsprings ^26^. In the context of neonatal overweight reported here, epigenetic modifications in genes involved in immune reactivity later in life, decreased generation of specific regulatory T cells, and dysbiosis of the microbiota, are all credible mechanisms of pathological imprinting that remain to be assessed.

Our study demonstrates that neonatal overweight is a risk factor for the development of colitis later in life. The multiple intestinal perturbations induced by excess calorie intake early in life, and their causal role in pathological imprinting, offer preventive measures that must be applied before and during weaning. It is also desirable to identify measures to reverse pathological imprinting after weaning, once the offspring has been recognized as having experienced such risk factors. The identification of the nature of pathological imprinting will offer potential avenues to reverse to healthy imprinting.

## ACKNOWLEDGMENTS

We thank all the members of the Microenvironment & Immunity Unit, as well as from the Stroma, Inflammation & Tissue Repair Unit, for support and discussion. We also thank the members of Gnotobiology Platform of the Institut Pasteur for technical support with germfree mice. Z.A.N. was supported by Pasteur-Roux Postdoctoral Fellowships from the Institut Pasteur. This work was supported by Institut Pasteur and INSERM, the Association François Aupetit, the Fondation pour la Recherche Medicale, and Innovator and Breakthrough awards from the Kenneth Rainin Foundation.

## AUTHOR CONTRIBUTIONS

Z.A.N. and G.E. planned the project and the experiments; Z.A.N., S.D., E.L., B.P. and M.B. performed the experiments, Z.A.N. analysed the data; M.B. provided germ-free resources; Z.A.N. and G.E. wrote the manuscript; Z.A.N. and G.E. acquired the funding; G.E. supervised the project.

## METHODS

### Mice

All animal experiments were approved by the committee on animal experimentation of the Institut Pasteur and by the French Ministry of Research. SPF females and males C57BL/6 mice were purchased from Charles River and exposed to the environment of the local animal facility at Institut Pasteur at least for 2 weeks before mating. Germ-free C57BL/6 mice were generated at the Gnotobiology Platform of the Institut Pasteur. All mice were weaned 28 days after birth (D28) and co-housing used in all experiments in order to avoid cage-specific divergence in the composition of the gut microbiota ^27^, experimental group were both age- and sex-matched. The age of mice at the beginning of each experiment is indicated in the legend to the figures.

### High fat diet and coconut oil gavage

Mice were fed HFD containing 60% of fat dominantly coconut oil (U8954 version 0140, SAFE, France) or normal chow diet (801954, SDS Diets) irradiated with 10kGys. Mice were supplemented with coconut oil (5 mL kg^−1^ day^−1^; 46949; Sigma; France) or a vehicle (V, distilled water) via oral gavage at 14 days until 28 days after birth.

### Control of litter size

Eight weeks old adult female C57BL/6 mice (Charles River, L’Arbresle, France) were individually housed. After two week of adaptation, they were mated overnight with males at a proportion of 2:1. The females were then housed in individual cages four days before delivery and during lactation. Mice were kept on a standard pellet diet (A03, SAFE Diets, France) under a 12-h light-dark cycle in a room maintained at a controlled temperature of 22°C and constant humidity. Litters were normalized to 7 pups per litter on postnatal day 1 (P1), with 4 male and 3 female pups per litter (NL = normal litter). On P3, some litters were culled to 3 pups per litter (2 male + 1 female pups, SL = small litter) to induce postnatal overfeeding. Animals were fed standard chow following weaning unless otherwise specified. Only male pups were used for these studies. Each experimental group in all experiments consisted of offsprings from at least 4 litters.

### Mice treatment with antibiotics, chemicals and antibodies

Mice were treated during the indicated periods of time with a cocktail of antibiotics in their drinking water, containing 2.5 mg/ml streptomycin, 1 mg/ml ampicillin, 0.5 mg/ml vancomycin, 1 mg/mL ciprofloxacin, and 0.5 mg/ml metronidazole. The antibiotic-containing drinking water was changed twice a week. Once the treatment was terminated, treated and untreated mice were co-housed. Where indicated, mice were injected i.p. 3x/week with 50μL of 5-ASA (5-aminosalicylic acid; 200μg/mL) to reduce H_2_S and sulfide reducing bacteria levels, or gavaged with 50 mg/kg of GYY4137 (Morpholin-4-ium (4-methoxyphenyl)morpholin-4-ylphosphinodithioate) or 10 μmol/kg NaHS. For MLCK inhibition, mice were treated by i.p. injection of ML-7 (2 mg/kg) 3x/week between 2 to 4 weeks after birth ^28^. For TNF-α and IFN-γ neutralisation, mice were treated simultaneously with a combination of 25 μg of anti-TNF-α mAb (clone MP6-XT22) and 25 μg of anti-IFN-γ mAb (XMG1.2) via i.p. injection 3x/week between 2 to 4 weeks after birth.

### Models of colitis

To test susceptibility to DSS-induced colitis, adult mice with similar body weight were exposed to two cycles of 2.5% DSS (with a mol. wt. of approx. 40.000 g/mol) in their drinking water (except GF mice that received 1% of DSS), for 7 days interrupted by 7 days of normal water. Control mice received drinking water without DSS. Progression of disease was monitored daily by weight and survival. Colitis severity was scored by daily observation of the following parameters: weight loss (0 point = No weight loss or weight gain, 1 point = 5–10% weight loss, 2 points = 11-15% weight loss, 3 points = 16-20% weight loss, 4 points = >21% weight loss); stool consistency (0 point = normal and well formed, 2 points = very soft and unformed, 4 points = watery stool); and blood in stool (0 point = normal color stool, 2 points = reddish color stool, 4 points = bloody stool). The disease activity index (DAI) was calculated as the combined scores of weight loss, bleeding, stool consistency, with a maximum score of 12. After 21 days, mice were euthanized, the length of colons was measured, and organs and blood were collected for biochemical analysis and a small piece (0.2cm) of distal colon was taken for the analysis of gene expression.

### Gut permeability

*In vivo* permeability assays were performed using fluorescein isothiocyanate (FITC)–dextran 4kDa as a paracellular permeability tracer. FITC–dextran was given by gavage to mice 6 days after initiation of the second cycle of DSS in the DSS-induced colitis model. Mice were gavaged with FITC–dextran (5 mg/200 μL/mouse) 4 h prior to sacrifice. Whole blood FITC–dextran concentration was determined by spectrometry. FITC–dextran concentrations in serum were calculated from standard curves generated by serial dilution of FITC–dextran. *In vivo* permeability was expressed as the mean whole blood FITC–dextran concentration in ng/mL.

### ELISA

Myeloperoxidase (MPO) and Lipocalin-2 were measured using ELISAs and following the manufacturers’ recommendations (from R&D Systems).

### Measure of H_2_S

Fecal and colonic content H_2_S levels were quantified using the methylene blue method ^29^. Colonic contents were collected after animals were euthanized. Material (0.1g) was homogenized with 1% zinc acetate trapping solution, 20 mM N,N-dimethyl-p-phenylenediamine sulfate prepared in 7.2 N HCl, and 30 mM FeCL3 prepared in 1.2 N HCl, and incubated for 30 minutes. Samples were centrifuged, and the clear upper phase was analyzed at 670 nm in comparison with a calibration curve of standard H_2_S solutions ^30^. H_2_S levels were expressed in nM/g weight.

### Quantification of sulfite-reducing bacteria

Fecal genomic DNA was extracted from the weighted stool samples as previously described ^31^, a method based on Godon DNA extraction. Sulfite-reducing bacteria were quantified using specific primers for the *dsrA* gene (encoding a dissimilatory sulfite reductase alpha subunit) as described by Devkota ^10^: F: 5’-CCA ACATGCACGGYTCCA-3”, R: 5’-CGTCGA ACTTGA ACTTGA ACTTGT AGG-3’.

### Quantitative PCR

Frozen tissue samples were dissociated in lysing/binding buffer of Multi-MACS cDNA kit (Miltenyi) with 0.5% antifoam using the gentleMACS™ Octo Dissociator (Miltenyi). RNA isolation, cDNA synthesis and cDNA purification were performed using the MultiMACS M96thermo Separation Unit following the manufacturer’s instructions of the MultiMACS ™ cDNA Synthesis Kit (Miltenyi). Real-time quantitative PCR cDNA was performed using SybrGreen (BioRad) and Qiagen primers. Ct values were normalised to the mean Ct obtained with the three housekeeping genes, Gapdh, Hsp90 and Hprt, in each sample.

### Statistical analysis

Statistical analysis was performed in GraphPad Prism statistical software (version 7). *P* values were calculated using the test indicated in the legends to the figures.

## LEGEND TO SUPPLEMENTARY FIGURES

**Supplementary Figure 1** Overweight induced by excessive calorie intake early in life did not persist into adulthood. **(a** and **d)** Mice exposed to high fat diet (HFD) until 4 weeks of age were weaned to normal chow (NC). **(b** and **e)** Mice fed NC were supplemented or not with coconut oil between 2 to 4 weeks of age. **(c** and **f)** The number of mice reared to NC was reduced at day 3 post-birth to 3 pups (SL; small litter) or maintained to 7 pups per litter as normal litter (NL). Weight measure of total body, epididymal fat, retroperitoneal fat and liver weights **(a-c)** at 4 weeks of age or **(d-f)** before DSS induction in adulthood. For **a** (n=30/group), **b** and **c** (n=8/group), **d** and **f** (n=10/group) and **e** (n=6-8/group). Mann Whitney test (\*\*\**P* < 0.001 and \*\*\*\**P* < 0.0001). ns = not significant. Data were pooled from at least three independent experiments. Data are shown as mean ± s.e.m.

**Supplementary Figure 2** Exposure to HFD early in life, but not after, determines colitis severity in adult mice. **(a)** Percentage of survival, **(b)** diseases activity index, **(c)** level of myeloperoxidase (MPO), **(d)** intestinal permeability and **(e)** colonic transcript profile in adult mice exposed to normal chow (NC) or to high fat diet (HFD) until 4 weeks of age (early in life) or between 6 to 12 weeks after birth (adult age). NC group, **a** (n=8), **b** (n=7), **c-e** (n=15); HFD (early in life) group, **a** (n=13), b (n=5), **c-e** (n=14); HFD (at adult age) group, **a** (n=13), **b** (n=10), **c-e** (n=11); No colitis group, **c-e** (n=10). For **a**, Log-rank (Mantel–Cox) test (*P* = 0.009); **b-e**, One-way ANOVA with Sidak’s multiple comparisons test (\*\**P* < 0.01 and \*\*\**P* < 0.001). Data were pooled from at least three independent experiments. Data are shown as mean ± s.e.m.

**Supplementary Figure 3** Exposure to coconut oil during weaning determines colitis severity in adult mice. **(a)** Percentage of survival, **(b)** diseases activity index, **(c)** level of myeloperoxidase (MPO), **(d)** intestinal permeability and **(e)** colonic transcript profile in adult mice supplemented or not with coconut oil during 2 to 4 weeks after birth. Vehicle group, **a** (n=9), **b** (n=8), **c-d** (n=5), **e** (n=8); Coconut oil group, **a** (n=19), **b** (n=9), **c-d** (n=5), e (n=8); No colitis group, **c-d** (n=3). For **a**, Log-rank (Mantel–Cox) test (*P* = 0.04); **b-e**, One-way ANOVA with Sidak’s multiple comparisons test (\**P* < 0.05; \*\**P* < 0.01; \*\*\**P* < 0.001 and \*\*\*\**P* < 0.0001). Data were pooled from at least three independent experiments. Data are shown as mean ± s.e.m.

**Supplementary Figure 4** Neonatal overfeeding determines colitis severity in adult mice. **(a)** Percentage of survival, **(b)** diseases activity index, **(c)** level of myeloperoxidase (MPO), **(d)** intestinal permeability and **(e)** colonic transcript profile in adult mice that were overfed by reducing litter size (SL; small litter) or not (NL; normal litter). NL group, **a-b** (n=8), **c** (n=4), **d** (n=6), **e** (n=8); SL group, **a** (n=14), b (n=8), **c** (n=4), **d-e** (n=8); No colitis group, **c-d** (n=4), **e** (n=5). For **a**, Log-rank (Mantel–Cox) test (*P* = 0.03); b-e, One-way ANOVA with Sidak’s multiple comparisons test (\**P* < 0.05; \*\**P* < 0.01; \*\*\**P* < 0.001 and \*\*\*\**P* < 0.0001). Data were pooled from at least three independent experiments. Data are shown as mean ± s.e.m.

**Supplementary Figure 5** Excessive calorie intake early in life did not impacts gut functionality before colitis induction in adult mice. **(a-c)** Colonic expression of mRNA coding for cytokines **(a-c, left)** or epithelial MLCK and tight junction proteins **(a-c, right)** and **(d-f)** intestinal permeability in adult mice exposed or not to antibiotics (ABX). **(a,d)** Mice were fed NC or HFD early in life, **(b,e)** supplemented or not with coconut oil during 2 to 4 weeks after birth, **(c,f)** overfeed by reducing litter size (SL; small litter) or not (NL; normal litter) at neonatal age. One-way ANOVA with Sidak’s multiple comparisons test (ns = not significant). **a,c,e** (n=6/group); **b** (n=4/group); **d** (n=8/group); **f** (n=10/group). Data were pooled from two independent experiments. Data are shown as mean ± s.e.m.

**Supplementary Figure 6** Excessive calorie intake early in life impacts intestinal gene expression. Colonic expression of mRNA for cytokines **(a,c,e)**, mucins and anti-microbial peptides **(b,d,f)** at 4 weeks after birth of mice exposed or not to antibiotics (ABX). **(a-b)** Mice were fed NC or HFD early in life, **(c-d)** supplemented or not with coconut oil during 2 to 4 weeks after birth, **(e-f)** overfeed by reducing litter size (SL; small litter) or not (NL; normal litter) at neonatal age. One-way ANOVA with Sidak’s multiple comparisons test (\**P* < 0.05; \*\**P* < 0.01; \*\*\**P* < 0.001 and \*\*\*\**P* < 0.0001). For **a-b** (n=3-4/group)**, c-d** (n=4-6/group)**, e-f** (n=3-4/group), the data is representative of one among two independent experiments. Data are shown as mean ± s.e.m.

**Supplementary Figure 7** Excessive calorie intake increases body weight through microbiota-dependent mechanism. Weight of total body **(a,e,i)**, epididymal fat **(b,f,j)**, retroperitoneal fat **(c,g,k)** and liver **(d,h,l)** at 4 weeks after birth of mice exposed or not to antibiotics (ABX). Mice were **(a-d)** fed NC or HFD early in life, **(e-h)** supplemented or not with coconut oil during 2 to 4 weeks after birth, **(i-l)** overfeed by reducing litter size (SL; small litter) or not (NL; normal litter) at neonatal age in SPF or in GF conditions. One-way ANOVA with Sidak’s multiple comparisons test (\**P* < 0.05; \*\**P* < 0.01; \*\*\**P* < 0.001 and \*\*\*\**P* < 0.0001). For **a-b** (n=3-4/group)**, c-d** (n=4-6/group)**, e-f** (n=3-4/group), the data is representative of one among two independent experiments. Data are shown as mean ± s.e.m.

**Supplementary Figure 8** Neutralization of pro-inflammatory cytokines reduced gut abnormalities in mice fed HFD early in life. **(a-c)** Weight of **(a)** epididymal fat, **(b)** retroperitoneal fat, **(c)** liver and **(d)** expression of mRNA for mucins and antimicrobial peptides (top) and cytokines (bottom) at 4 weeks of age in mice fed NC or HFD, and treated with anti-IFNγ and anti-TNFα (anti-I+T) neutralizing antibodies (mAb) or isotypes controls during 2 to 4 weeks of age. One-way ANOVA with Sidak’s multiple comparisons test (\**P* < 0.05; \*\**P* < 0.01; \*\*\**P* < 0.001 and \*\*\*\**P* < 0.0001). For **a,b,c** (n=8/group), the data were pooled from at least two independent experiments. For **d** (n=5-6/group), the data is representative of one among two independent experiments. Data are shown as mean ± s.e.m.

**Supplementary Figure 9** Neutralization of pro-inflammatory cytokines induced by HFD early in life reduced colitis severity in adult mice. **(a)** Percentage of survival, **(b)** disease activity index, **(c)** level of myeloperoxidase (MPO), **(d)** intestinal permeability and **(e)** colonic transcript profile in adult mice exposed to NC or to HFD early in life and treated with anti-IFNγ and anti-TNFα (anti-I+T) neutralizing antibodies (mAb) or isotypes controls during 2 to 4 weeks of age. For **a**, Log-rank (Mantel–Cox) test (*P* = 0.009); **b-e**, One-way ANOVA with Sidak’s multiple comparisons test (\**P* < 0.05; **P < 0.01; \*\*\**P* < 0.001 and \*\*\*\**P* < 0.0001). ns = not significant. NC+isotype mAb group, **a** (n=6), b (n=6), **c-e** (n=4); HFD+isotype mAb group, **a** (n=8), **b** (n=7), **c-e** (n=5); NC+anti-I+T mAb group, **a** (n=8), **b** (n=7), **c-e** (n=4); HFD+anti-I+T mAb group, **a** (n=14), **b** (n=6), **c-e** (n=5); No colitis group, **c-e** (n=4). Data were pooled from at least two independent experiments. Data are shown as mean ± s.e.m.

**Supplementary Figure 10** Inhibition of MLCK reduced gut abnormalities in mice fed HFD early in life. **(a-c)** Weight of **(a)** epididymal fat, **(b)** retroperitoneal fat, **(c)** liver and **(d)** expression of mRNA for mucins and antimicrobial peptides (top) and cytokines (bottom) at 4 weeks of age from mice fed NC or HFD and treated with ML-7 or vehicle controls during 2 to 4 weeks of age. One-way ANOVA with Sidak’s multiple comparisons test (\**P* < 0.05 and \*\*\*\**P* < 0.0001). ns = not significant. For **a,b,c**, NC+Vehicle (n=8); HFD+Vehicle (n=8); NC+ML-7 (n=6); HFD+ML-7 (n=10); the data were pooled from at least two independent experiments. For **d**, NC+Vehicle (n=4); HFD+Vehicle (n=3); NC+ML-7 (n=4); HFD+ML-7 (n=4); the data is representative of one among two independent experiments. Data are shown as mean ± s.e.m.

**Supplementary Figure 11** Inhibition of intestinal permeability induced by HFD early in life reduced colitis severity in adult mice. **(a)** Percentage of survival, **(b)** disease activity index, **(c)** level of myeloperoxidase (MPO), **(d)** intestinal permeability and **(e)** colonic transcript profile after colitis induction in adult mice exposed to NC or to HFD early in life and treated with ML-7 or vehicle controls during 2 to 4 weeks of age. For **a**, Log-rank (Mantel–Cox) test (*P* = 0.14); **b-e**, One-way ANOVA with Sidak’s multiple comparisons test (\**P* < 0.05; \*\**P* < 0.01; \*\*\**P* < 0.001 and \*\*\*\**P* < 0.0001). ns = not significant. NC+vehicle group, **a** (n=6), **b** (n=4), **c-e** (n=4); HFD+vehicle group, **a** (n=14), **b** (n=5), **c-e** (n=5); NC+ML-7 group, **a** (n=5), **b** (n=4), **c-e** (n=4); HFD+ML-7 group, **a** (n=6), **b** (n=5), **c-e** (n=5); No colitis group, **c-e** (n=4). Data were pooled from at least two independent experiments. Data are shown as mean ± s.e.m.

**Supplementary Figure 12** The expression of bacterial *dsrA* and H_2_S levels. **(a-b)** The expression of bacterial *dsrA* coding for the dissimilative sulfite reductase and **(b)** at 4 weeks after birth in mice grown in reduced litters (SL, Small Litter) or normal litters (NL). For **a** (n=10/group), **b** (n=34/group). **(c-h)** Levels of H_2_S in feces from mice grown in SL (n=43) or NL (n=34), **(d)** in feces from mice gavaged with coconut oil (n=16) or not (n=8), **(e)** in the colon of mice fed NC or HFD (n=12/group), **(f)** in the colon of mice grown in SL or NL (n=21/group), **(g)** in feces of mice fed HFD and treated or not with antibiotics (n=10), and **(h)** in feces of mice grown in SL and treated or not with antibiotics (n=19), all measured at 4 weeks of age. **(i)** The expression of bacterial *dsrA* and **(j)** levels of H_2_S in feces from mice fed HFD and treated or not with 5-ASA (n=15/group) at 4 weeks of age. Mann Whitney test (\*\**P* < 0.01, \*\*\**P* < 0.001 and \*\*\*\**P* < 0.0001). ns = not significant. Data are shown as mean ± s.e.m.

**Supplementary Figure 13** Administration of 5-ASA suppresses abnormalities induced in mice fed HFD early in life. **(a-c)** Weight of **(a)** epididymal fat, **(b)** retroperitoneal fat, **(c)** liver and **(d)** expression of mRNA for mucins and antimicrobial peptides (top) and cytokines (bottom) at 4 weeks of age in mice fed NC or HFD and treated with 5-ASA or vehicle controls during 2 to 4 weeks of age. One-way ANOVA with Sidak’s multiple comparisons test (\**P* < 0.05; \*\**P* < 0.01; \*\*\**P* < 0.001 and \*\*\*\**P* < 0.0001). For **a,b,c**, NC+Vehicle (n=8); HFD+Vehicle (n=8); NC+5-ASA (n=5); HFD+5-ASA (n=5); the data were pooled from at least two independent experiments. For **d**, NC+Vehicle (n=3); HFD+Vehicle (n=3); NC+5-ASA (n=6); HFD+5-ASA (n=6); the data is representative of one among two independent experiments. Data are shown as mean ± s.e.m.

**Supplementary Figure 14** Administration of 5-ASA in mice fed HFD early in life reduced colitis severity in adulthood. **(a)** Percentage of survival, **(b)** disease activity index, **(c)** colonic length, **(d)** levels of myeloperoxidase (MPO), **(e)** intestinal permeability, **(f)** levels of lipocalin-2 and **(g)** colonic transcript profile after colitis induction in adult mice exposed to HFD early in life and treated with 5-ASA or vehicle during 2 to 4 weeks of age. For **a**, Log-rank (Mantel–Cox) test (*P* = 0.03); **b-g**, One-way ANOVA with Sidak’s multiple comparisons test (\**P* < 0.05; \*\**P* < 0.01; \*\*\**P* < 0.001 and \*\*\*\**P* < 0.0001). ns = not significant. HFD+vehicle group, **a** (n=12), **b** (n=4), **c-e** (n=4); HFD+5-ASA group, **a** (n=12), **b** (n=4), **c-g** (n=4); No colitis group, **c-g** (n=3). Data were pooled from at least two independent experiments. Data are shown as mean ± s.e.m.

**Supplementary Figure 15** Abnormalities induced by excessive calorie intake are dependent on H_2_S. **(a-c)** Weight of **(a)** epididymal fat, **(b)** retroperitoneal fat, **(c)** liver and **(d)** colonic expression of mRNA for mucins and antimicrobial peptides (top) and cytokines (bottom) at 4 weeks of age from germfree mice grown in SL and exposed to sodium hydrogen sulfide (NaHS) and/or heat-killed microbiota (M), or to vehicle control during 2 to 4 weeks of age. One-way ANOVA with Sidak’s multiple comparisons test (\**P* < 0.05; \*\**P* < 0.01; \*\*\**P* < 0.001 and \*\*\*\**P* < 0.0001). For **a,b,c**, SL+Vehicle (n=6); SL+NaHS (n=4); SL+M (n=4); SL+M+NaHS (n=4); the data were pooled from at least two independent experiments. For **d**, (n=4/group); the data is representative of one among two independent experiments. Data are shown as mean ± s.e.m.

**Supplementary Figure 16** Involvement of H_2_S in pathological imprinting induced by excessive calorie intake early in life. **(a)** Percentage of survival, **(b)** disease activity index, **(c)** colonic lenght, **(d)** colonic transcript profile and **(e)** intestinal permeability after colitis induction from adult germfree mice that are exposed to sodium hydrogen sulfde (NaHS) and/or heat-killed microbiota (M) or to vehicle, and grown in SL, during 2 to 4 weeks of age. For **a**, Log-rank (Mantel–Cox) test (*P* = 0.04); **b-e**, One-way ANOVA with Sidak’s multiple comparisons test (\**P* < 0.05; \*\**P* < 0.01; \*\*\**P* < 0.001 and \*\*\*\**P* < 0.0001). ns = not significant. SL+vehicle group, **a** (n=5), **b-e** (n=4); SL+NaHS group, **a** (n=5), **b-e** (n=4); SL+M group, **a** (n=7), **b-e** (n=4); SL+M+NaHS group, **a** (n=10), **b-e** (n=4); No colitis group, **c-e** (n=6). Data were pooled from at least two independent experiments. Data are shown as mean ± s.e.m.

**Supplementary Figure 17** Abnormalities induced by excessive calorie intake are dependent of H_2_S. Weight of **(a)** total body, **(b)** epididymal fat, **(c)** retroperitoneal fat, **(d)** liver, **(e)** intestinal permeability and (**f)** colonic expression of mRNA for cytokines (left) and MLCK and epithelial tight junction proteins at 4 weeks of age in germfree mice grown in SL and exposed to slow release of H_2_S (GYY4137) and/or heat-killed microbiota (M), or to vehicle control during 2 to 4 weeks of age. One-way ANOVA with Sidak’s multiple comparisons test (\**P* < 0.05; \*\**P* < 0.01; \*\*\**P* < 0.001 and \*\*\*\**P* < 0.0001). For **a-e**, SL+Vehicle (n=3); SL+GYY4137 (n=3); SL+M (n=4); SL+M+GYY4137 (n=5); the data were pooled from two independent experiments. For **f**, (n=3/group); the data is representative of one among two independent experiments. Data are shown as mean ± s.e.m.

**Supplementary Figure 18** Involvement of H_2_S in pathological imprinting induced by excessive calorie intake early in life. **(a)** Percentage of body loss, **(b)** disease activity index, **(c)** colonic length and **(d)** colonic transcript profile after colitis induction in adult germfree mice exposed to a slow-release H_2_S donor (GYY4137) and/or heat-killed microbiota (M) or to vehicle control, and grown in SL, during 2 to 4 weeks of age. One-way ANOVA with Sidak’s multiple comparisons test (\**P* < 0.05; \*\**P* < 0.01; \*\*\**P* < 0.001 and \*\*\*\**P* < 0.0001). ns = not significant. For **a-d** (n=3/group), the data is representative of one among two independent experiments. Data are shown as mean ± s.e.m.

**Supplementary Figure 19** Abnormalities induced by excessive calorie intake are dependent of H_2_S. Weight of **(a)** total body, **(b)** epididymal fat, **(c)** retroperitoneal fat, **(d)** liver, **(e)** intestinal permeability at 4 weeks of age in germfree mice grown in SL and exposed to sodium hydrogen sulfde (NaHS) and heat-killed microbiota (M), and treated with ML-7, with anti-IFNγ and anti-TNFα (anti-IT) neutralizing antibodies (mAb), or with vehicle control, during 2 to 4 weeks of age. **(f)** Percentage of body loss, **(g)** diseases activity index, **(h)** colonic length and **(i)** colonic transcript profile after colitis induction in adult germfree mice exposed to sodium hydrogen sulfde (NaHS) and heat-killed microbiota (M), and treated with ML-7, with anti-IFNγ and anti-TNFα (anti-IT) neutralizing antibodies (mAb) or with vehicle control, during 2 to 4 weeks of age. One-way ANOVA with Sidak’s multiple comparisons test (\**P* < 0.05; \*\*\**P* < 0.001 and \*\*\*\**P* < 0.0001). For **a-i**, No colitis group, SL group or SL+M+NaHS group (n=3); SL+M+NaHS+ML-7 or SL+M+NaHS+anti-IT group (n=4); the data is representative of one among two independent experiments. Data are shown as mean ± s.e.m.

