## Supplemental Figures for "Excess calorie intake early in life increases susceptibility to colitis in the adult"

**Supplementary Figure 1**

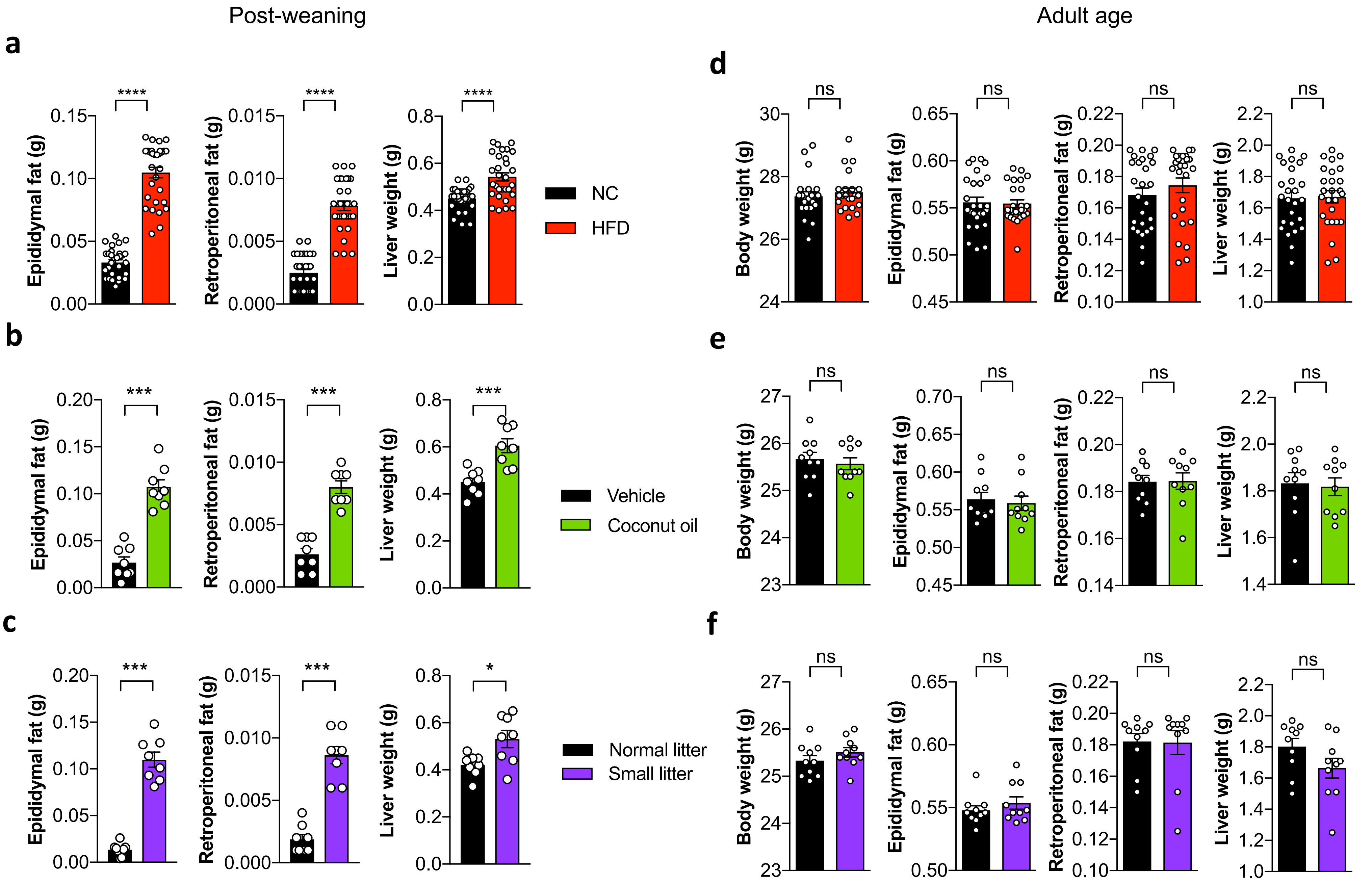

#### Supplementary Figure 2

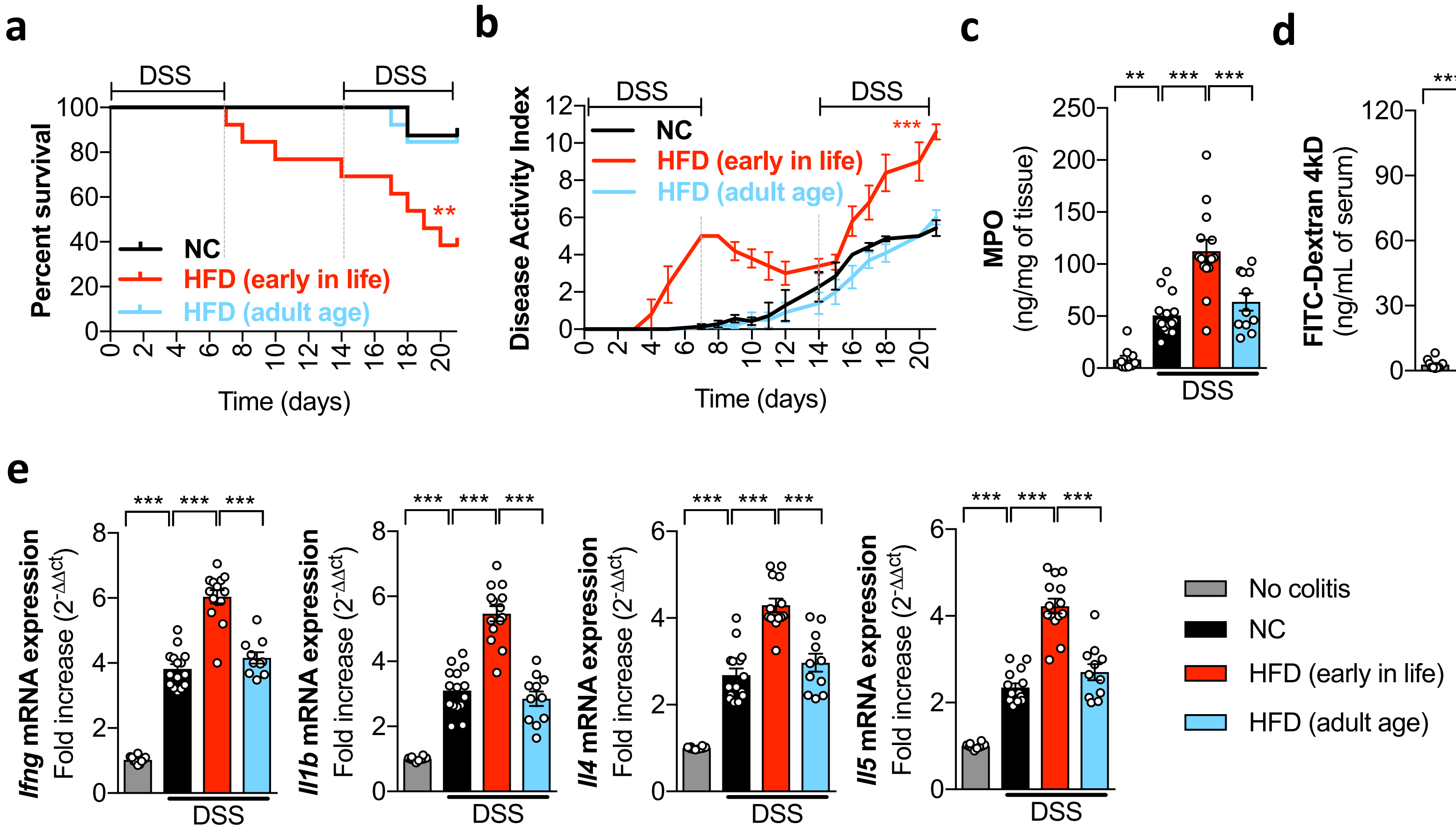

**Supplementary Figure 3**

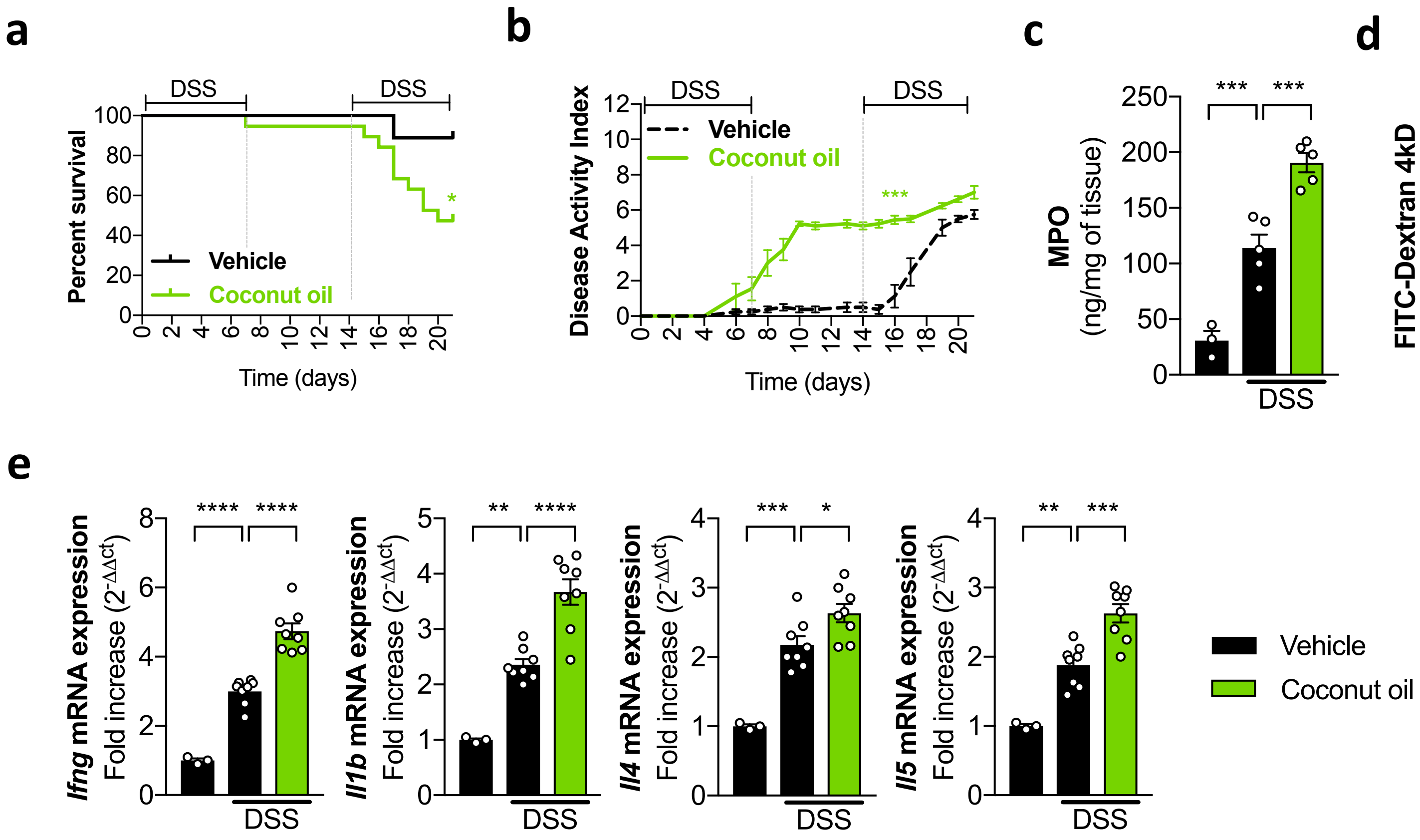

**Supplementary Figure 4**

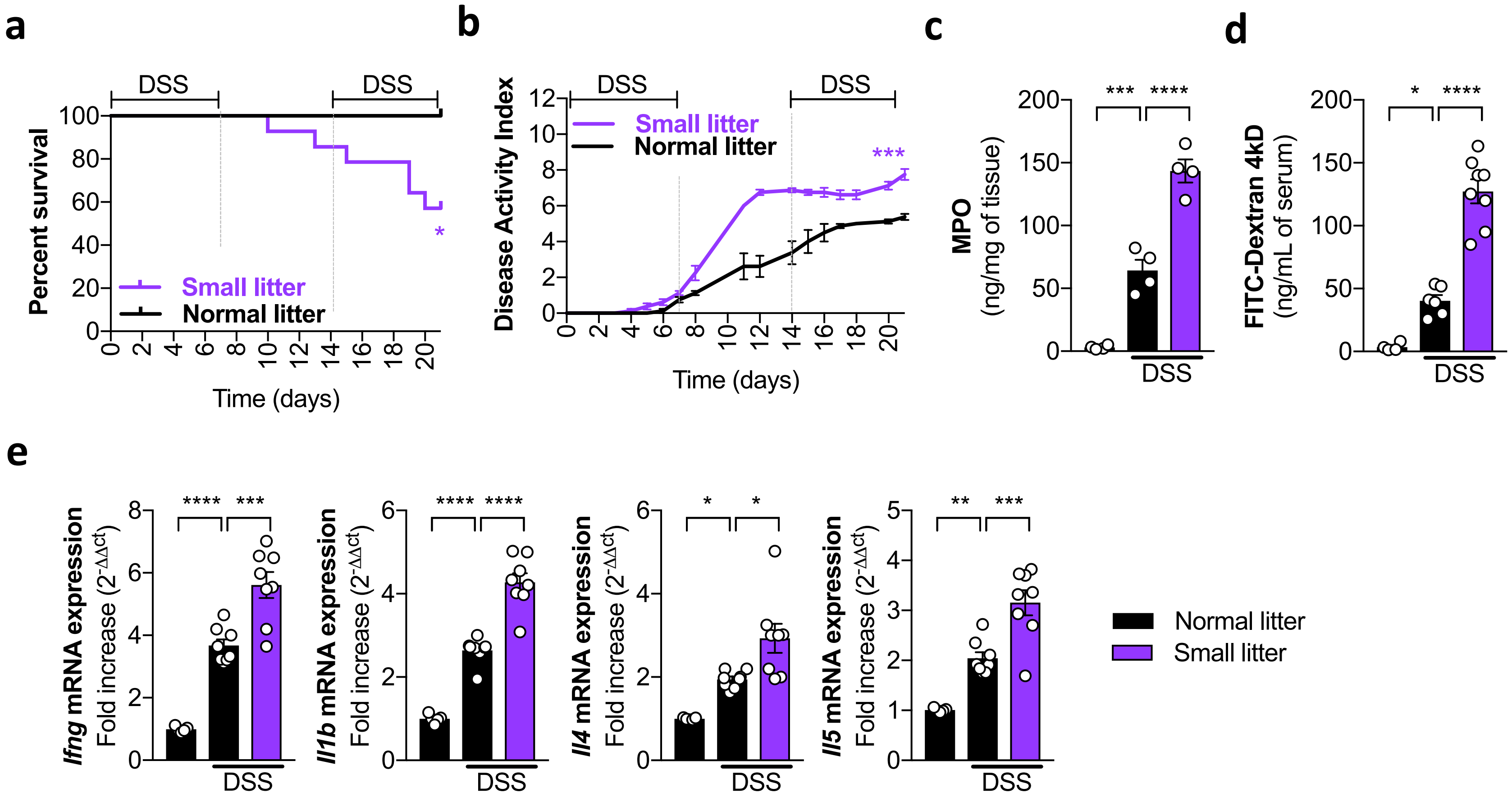

**Supplementary Figure 5**

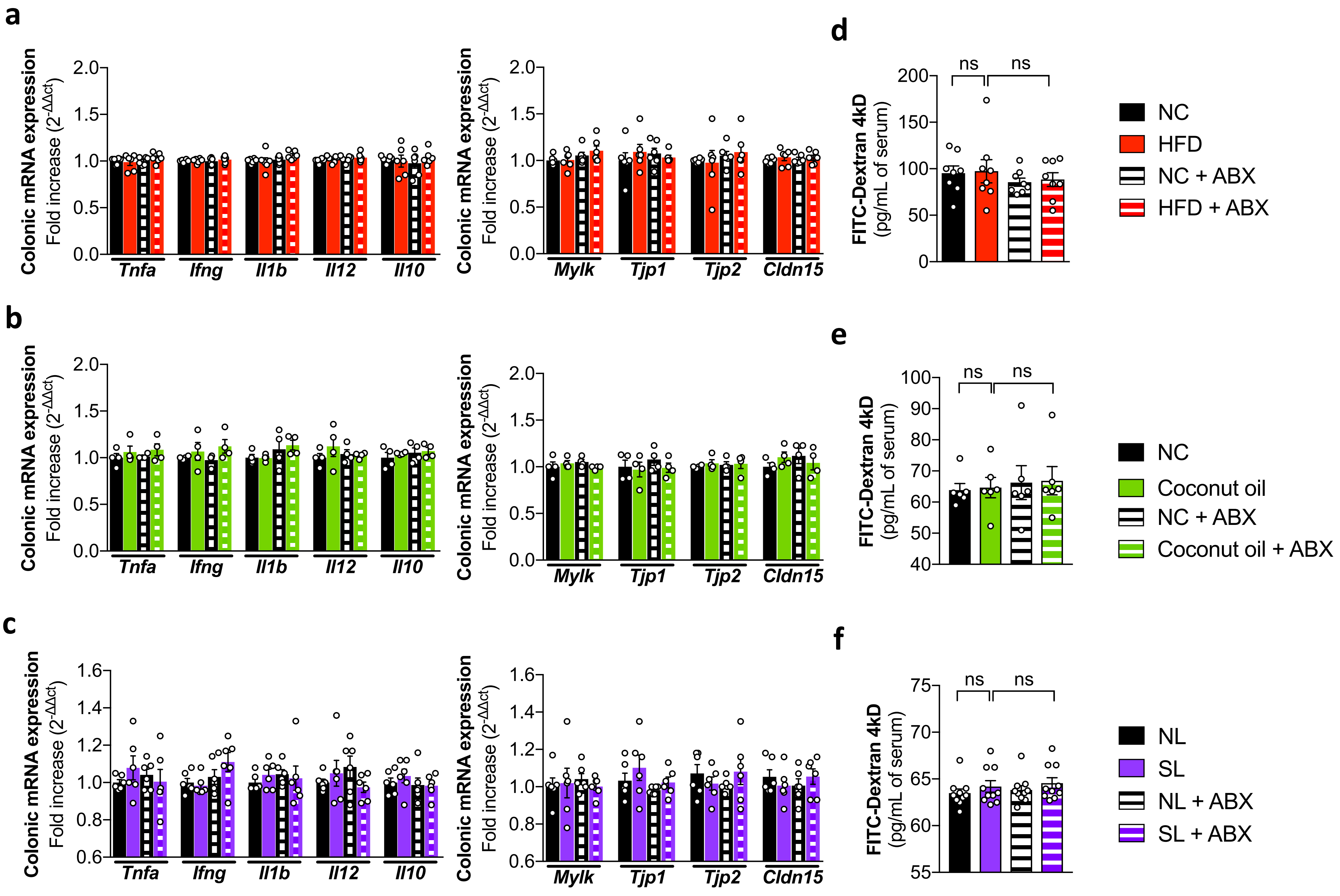

#### Supplementary Figure 6

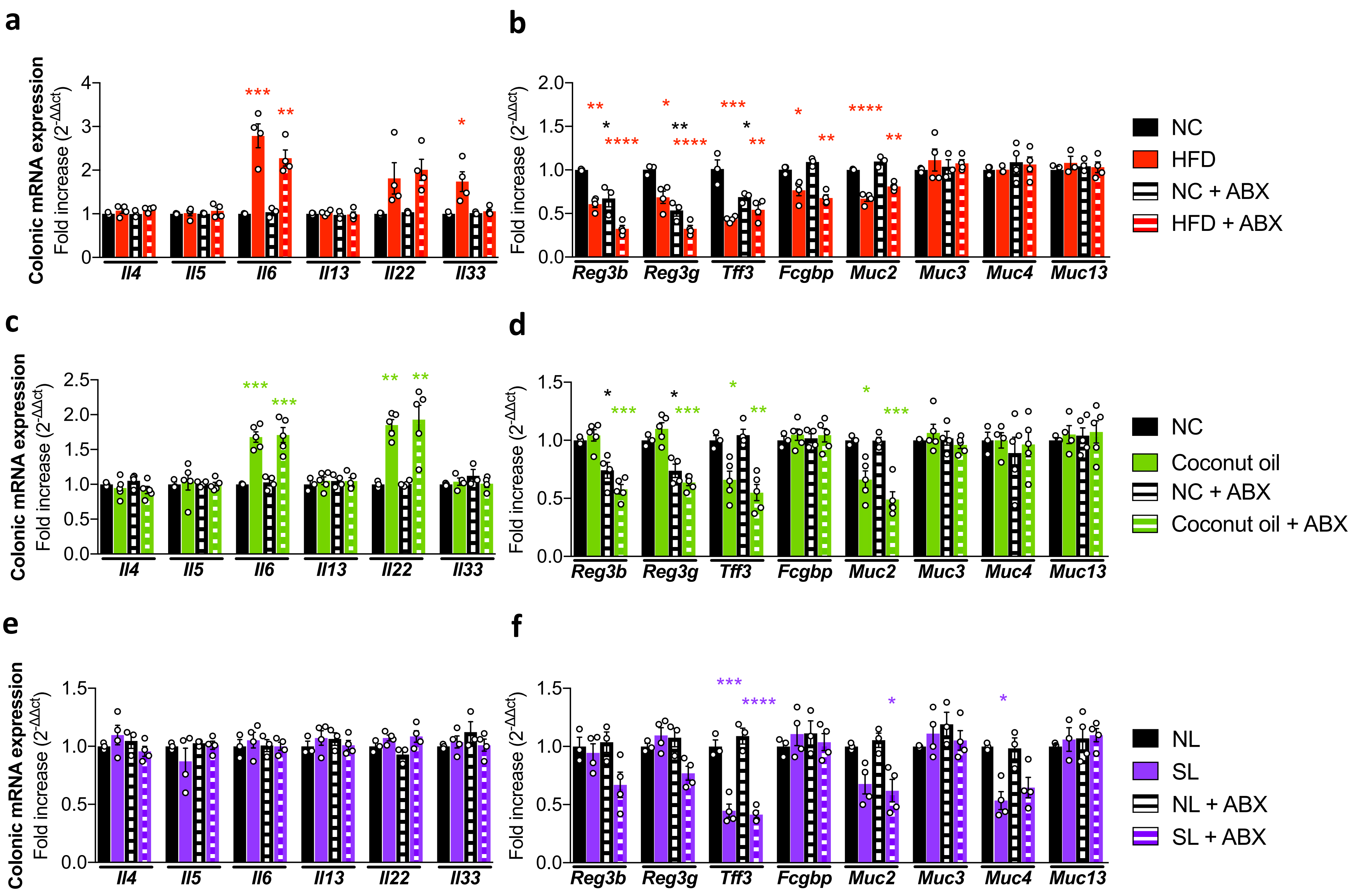

**Supplementary Figure 7**

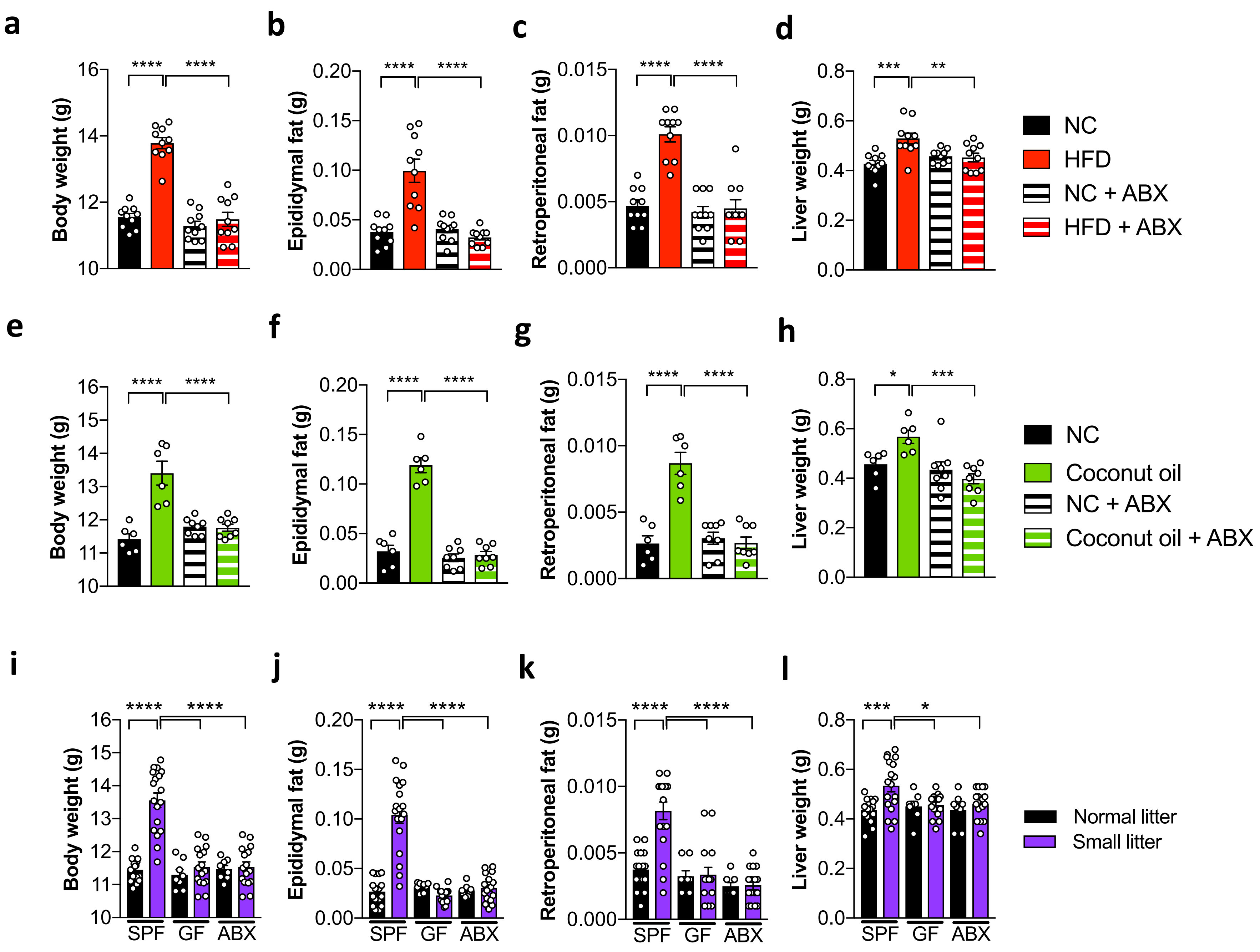

#### Supplementary Figure 8

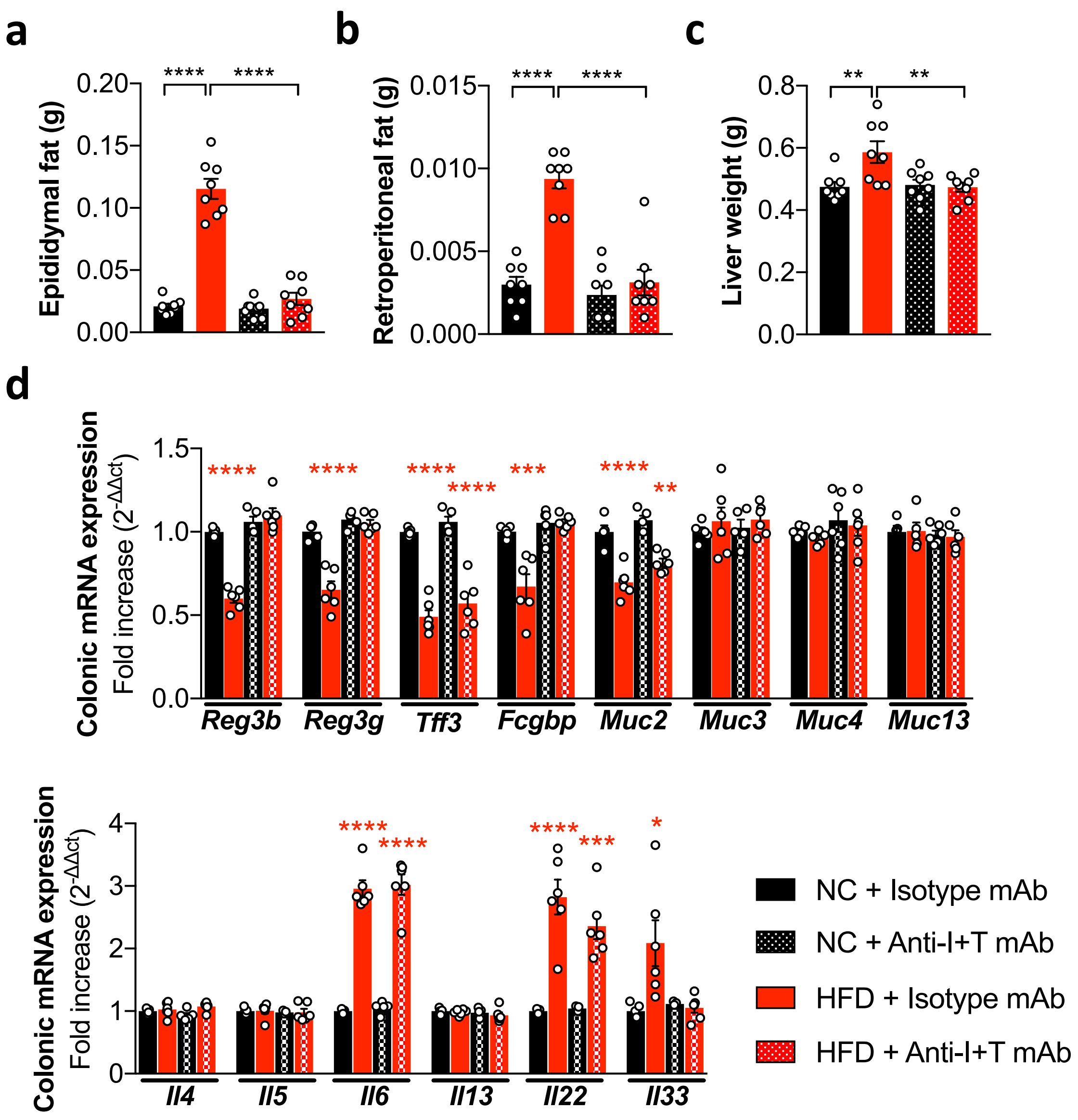

#### Supplementary Figure 9

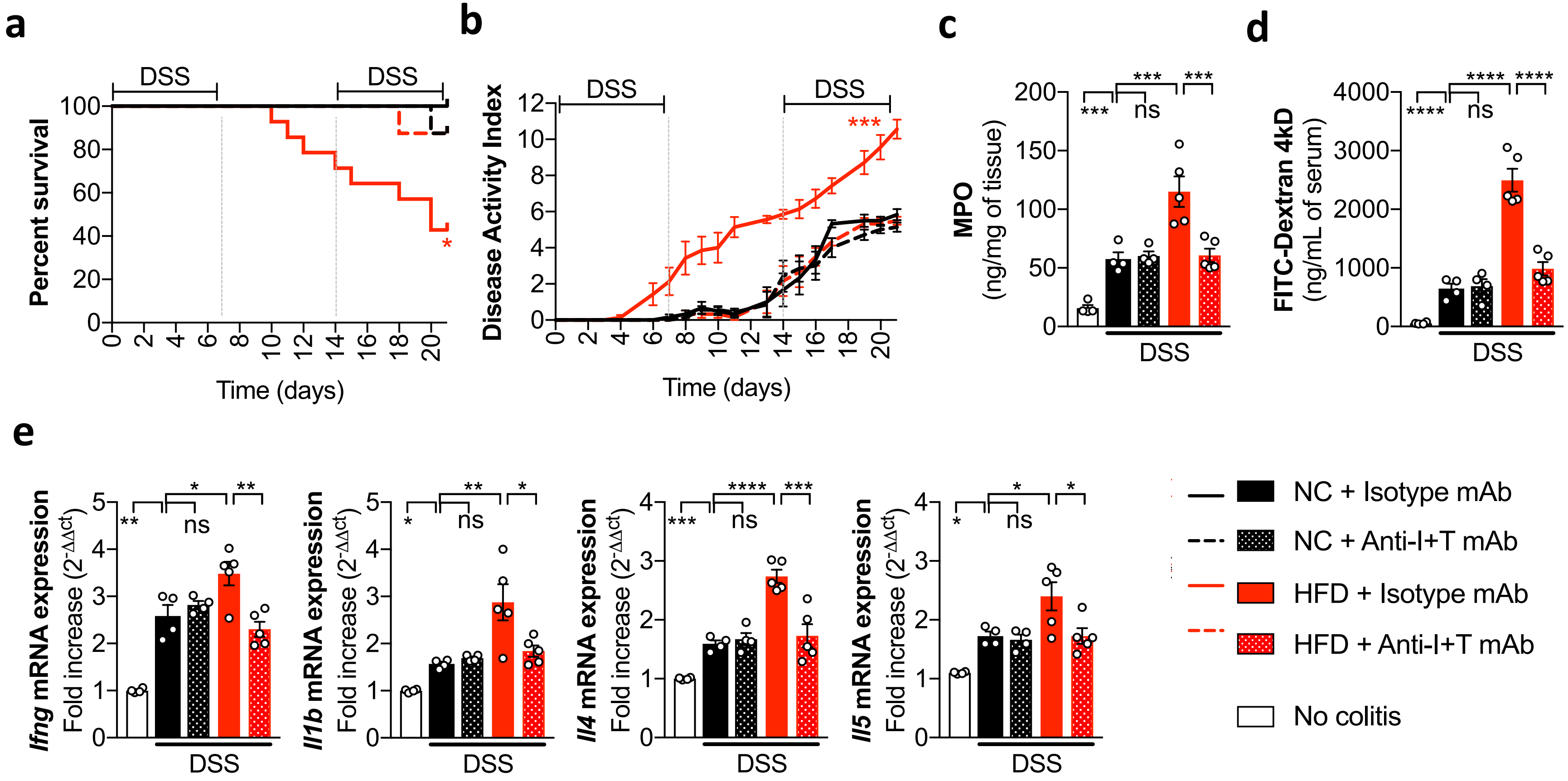

### Supplementary Figure 10

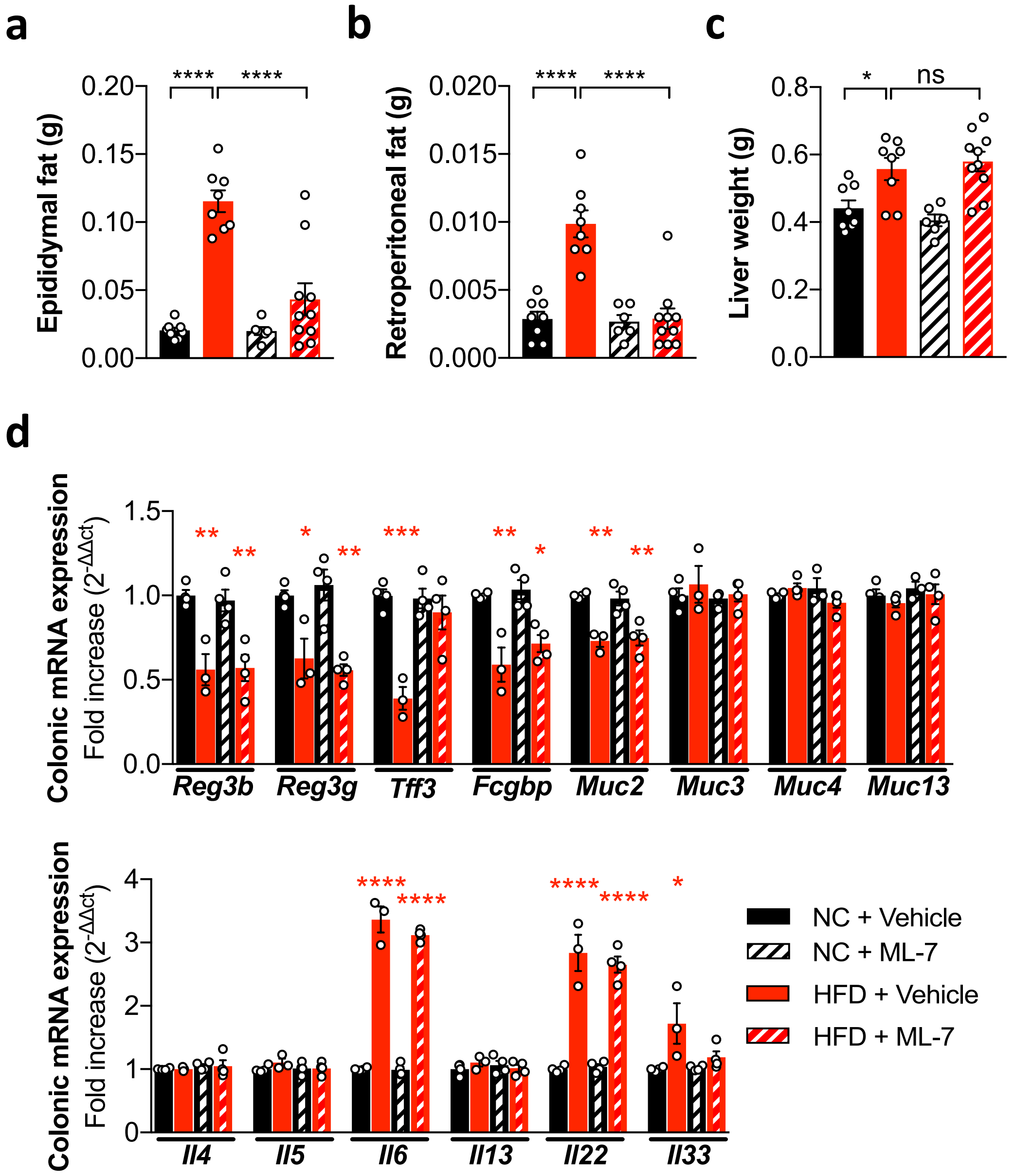

#### Supplementary Figure 11

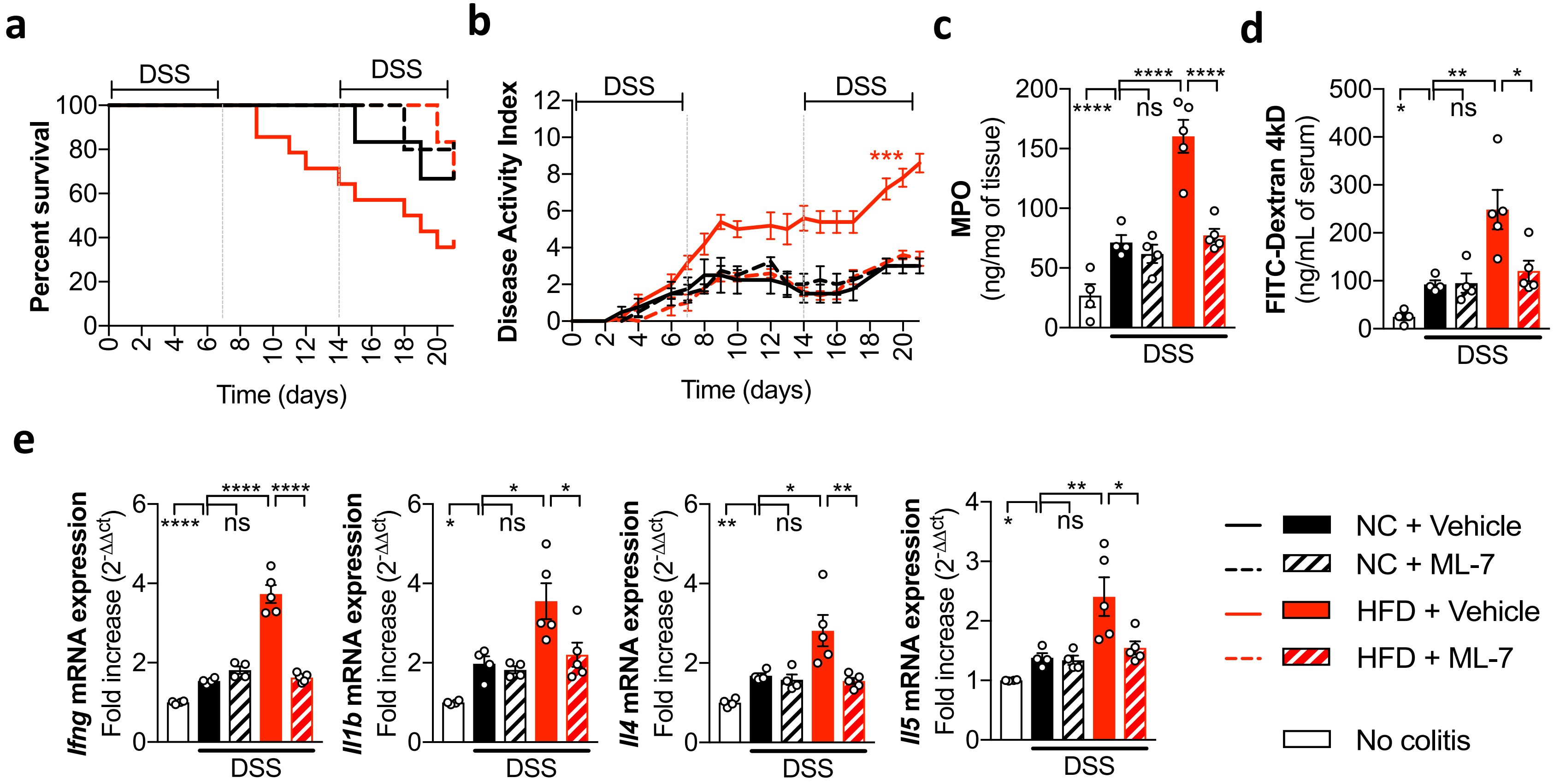

#### Supplementary Figure 12

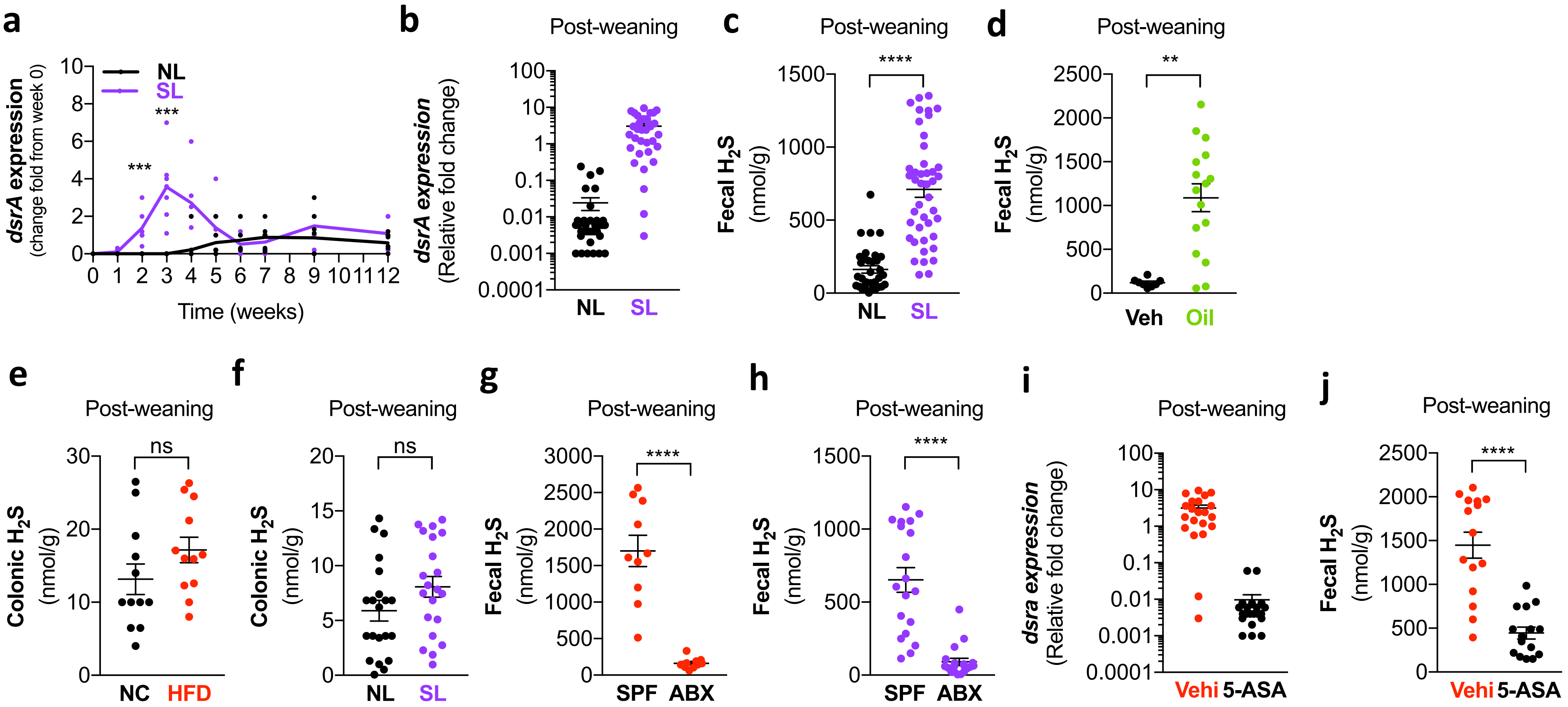

#### Supplementary Figure 13

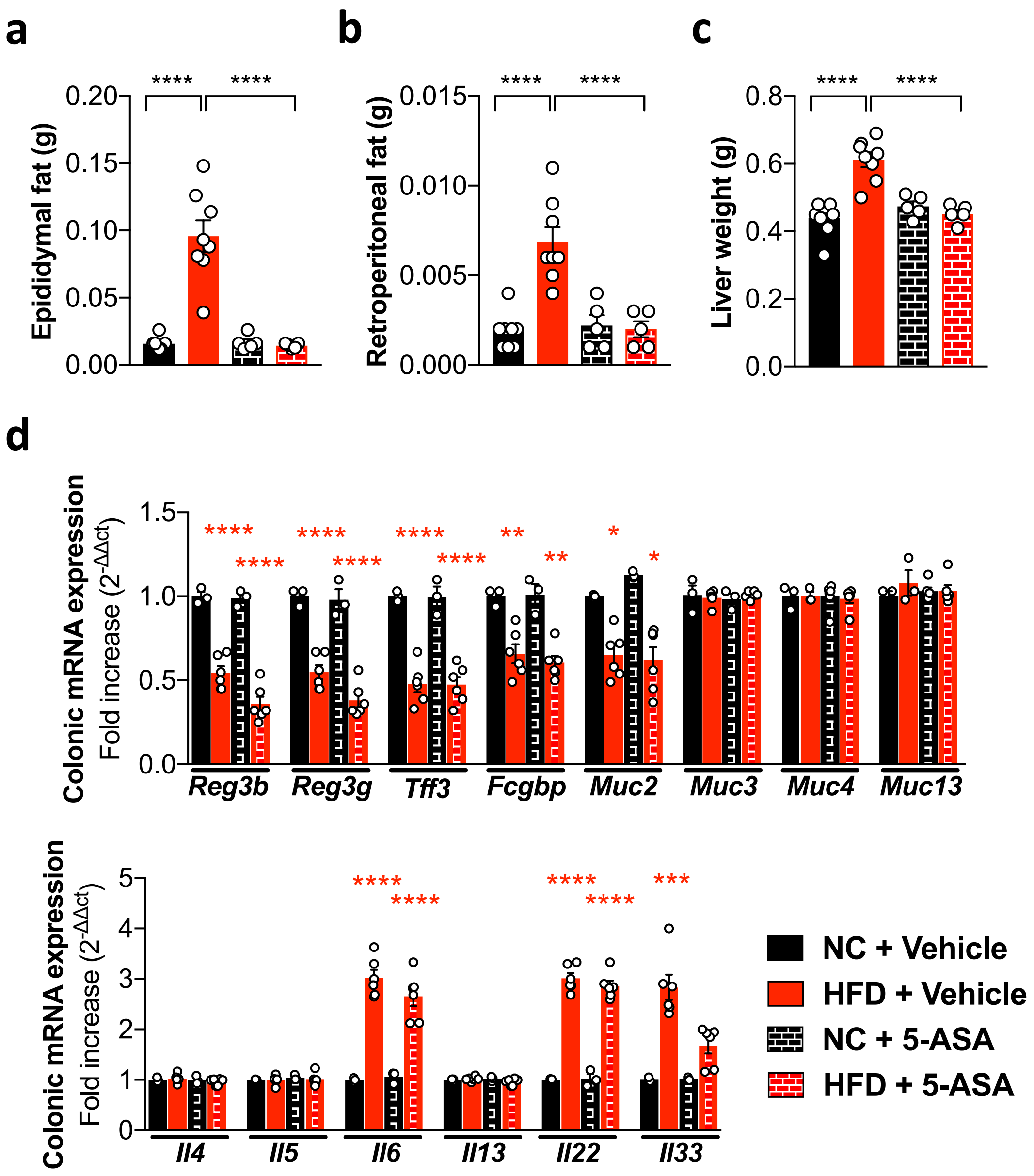

**Supplementary Figure 14**

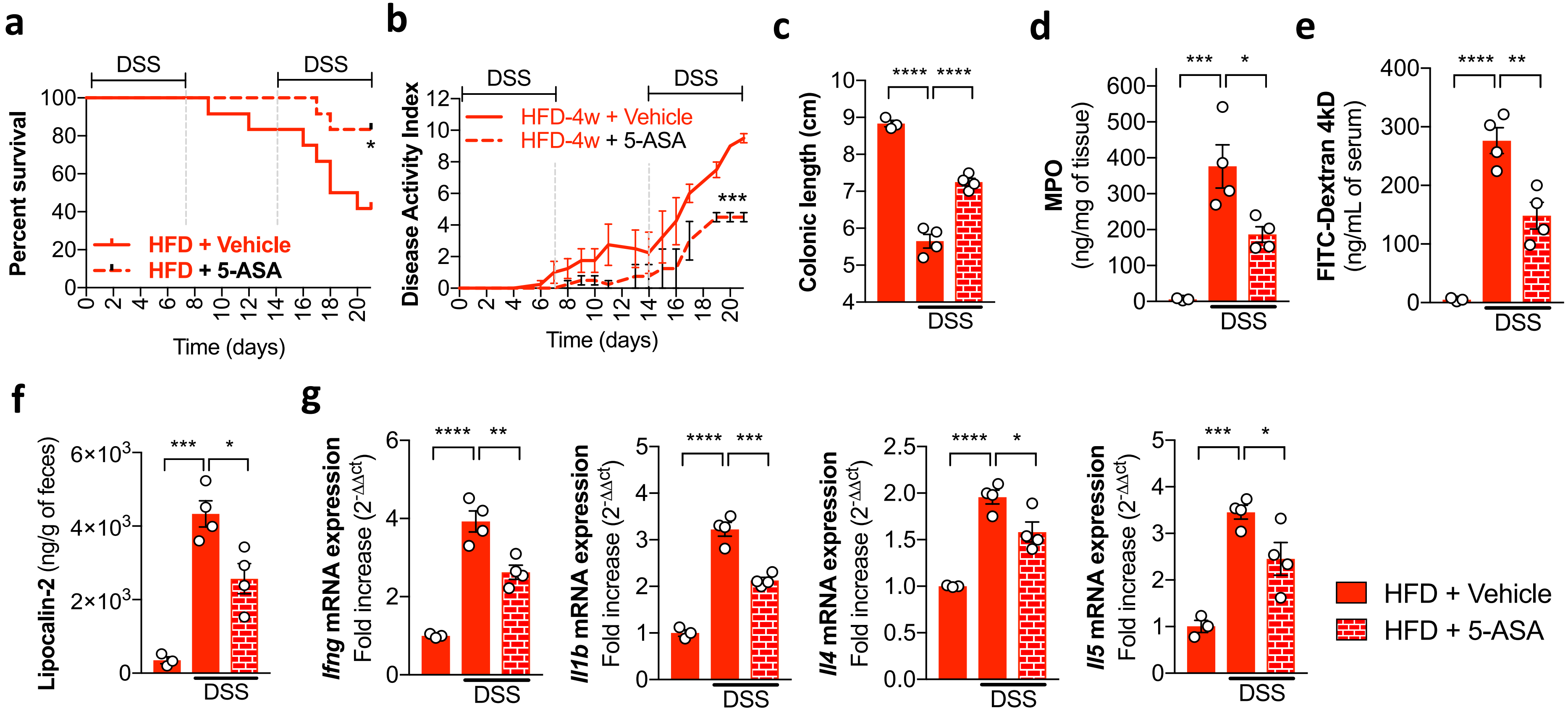

#### Supplementary Figure 15

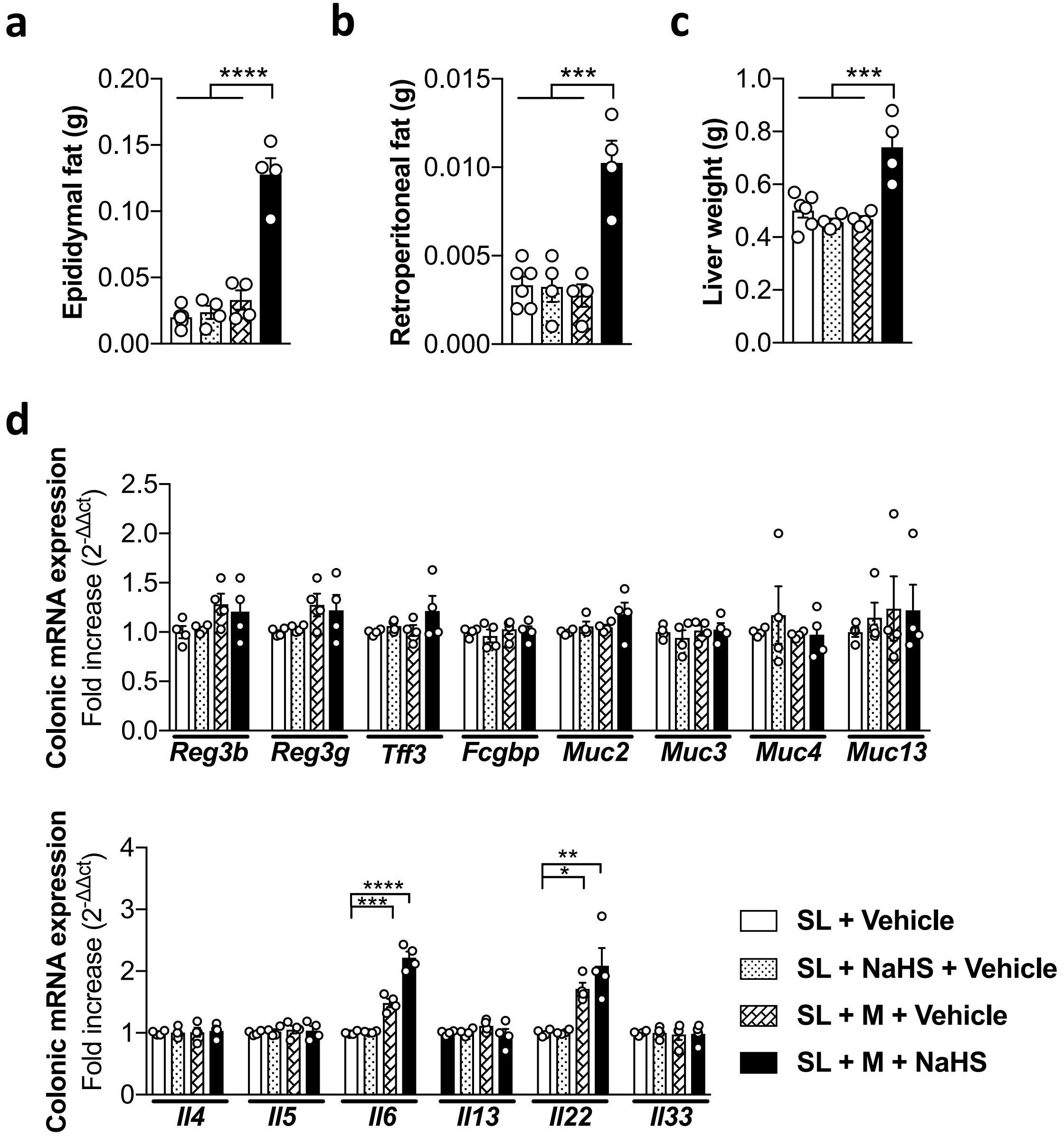

**Supplementary Figure 16**

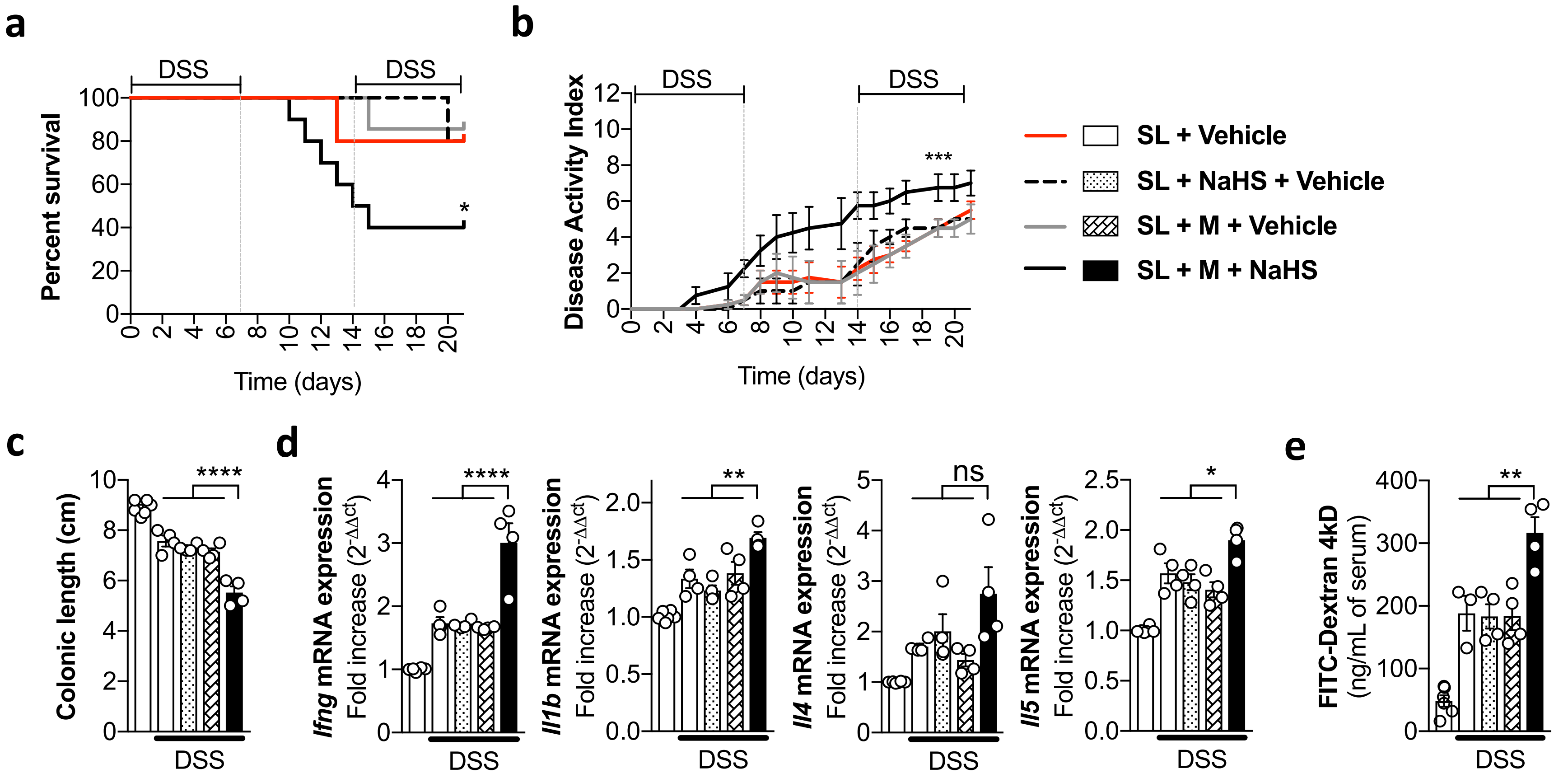

**Supplementary Figure 17**

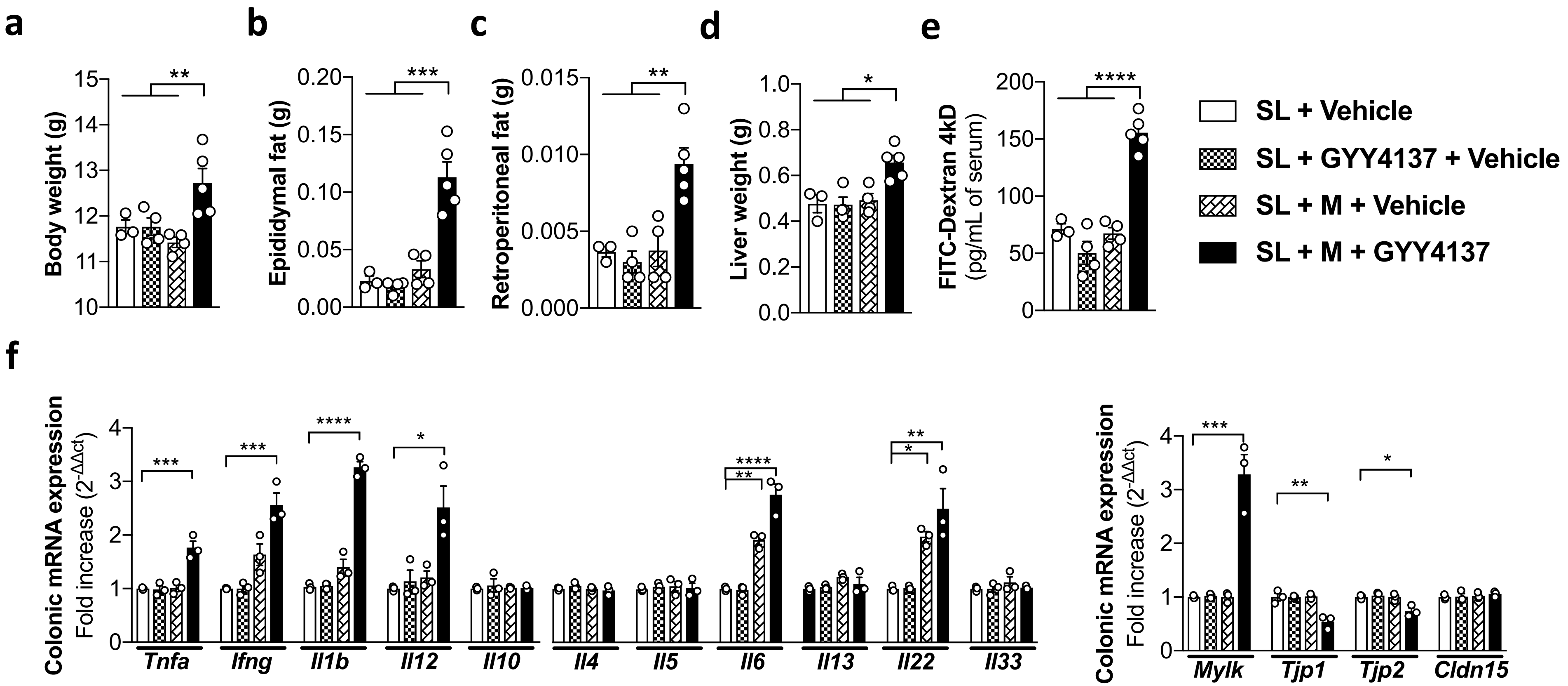

#### Supplementary Figure 18

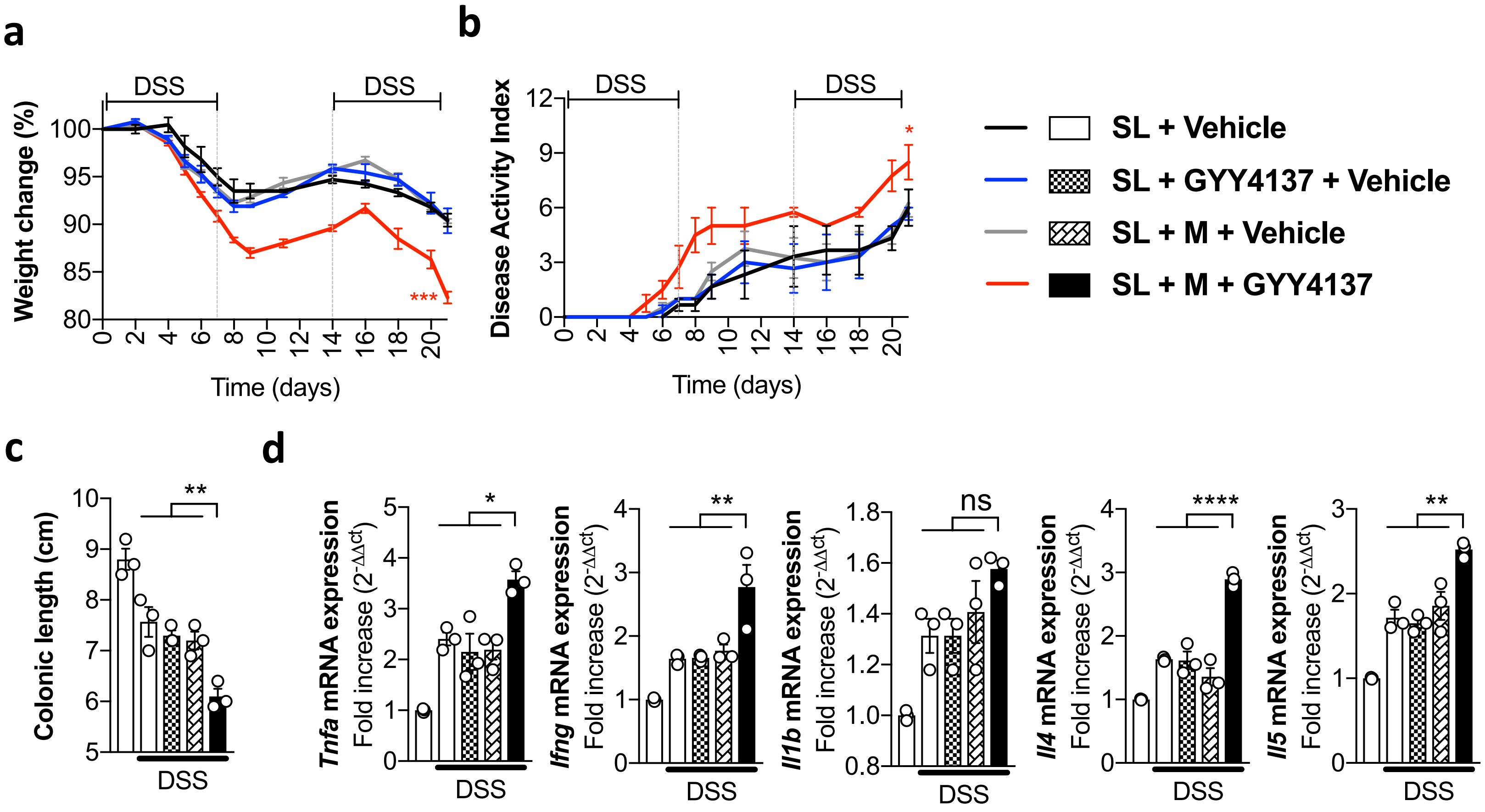

**Supplementary Figure 19**

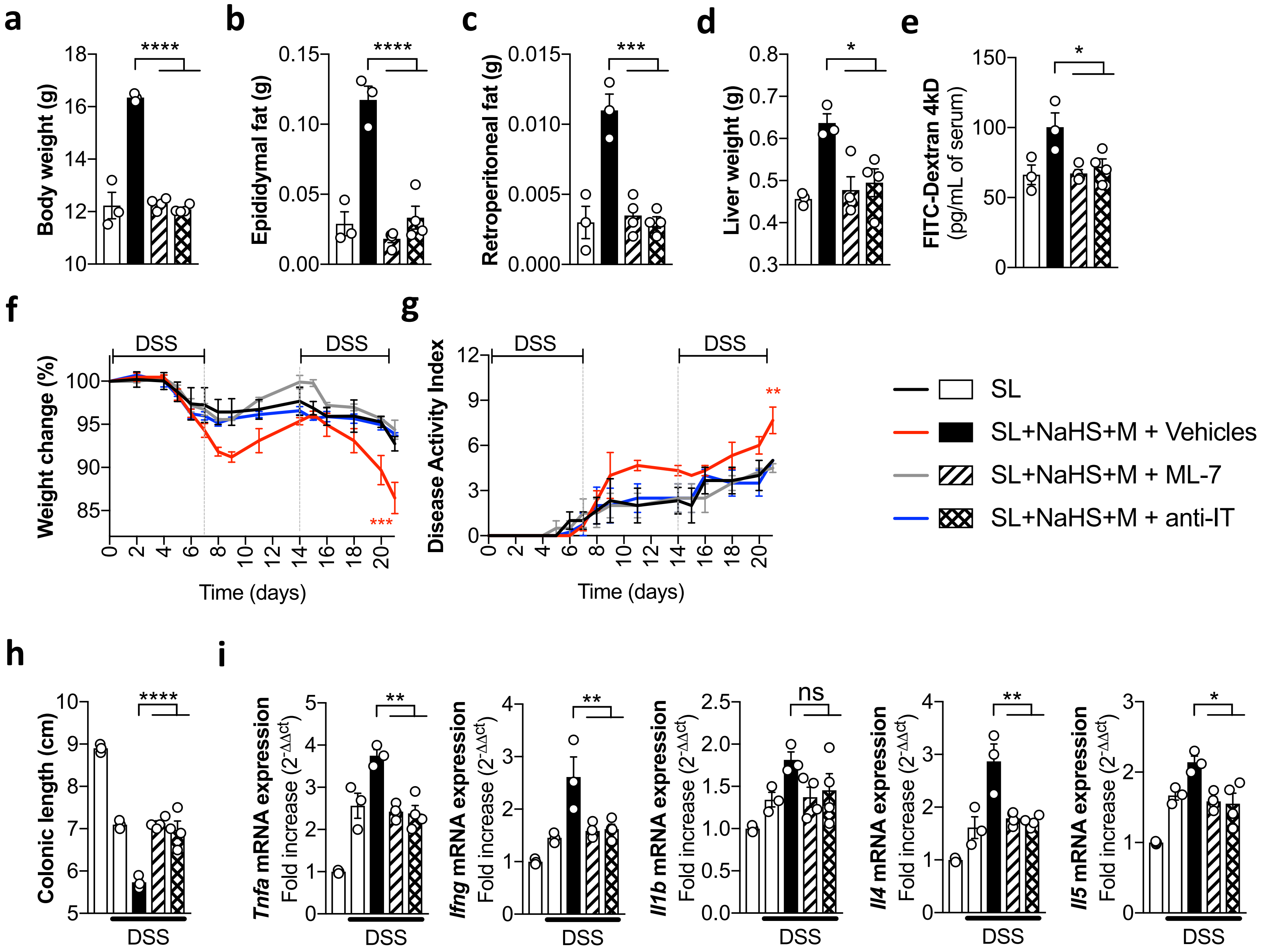
